# Glutamine addiction is targetable via altering splicing of nutrient sensors and epitranscriptome regulators

**DOI:** 10.1101/2024.02.28.582087

**Authors:** Jodie Bojko, Madhu Kollareddy, Marianna Szemes, Jacob Bellamy, Evon Poon, Ahmad Moukachar, Danny Legge, Emma E Vincent, Nicholas Jones, Sally Malik, Alex Greenhough, Alex Paterson, Ji Hyun Park, Kelli Gallacher, Louis Chesler, Karim Malik

## Abstract

About 50% of poor prognosis neuroblastoma arises due to MYCN over-expression. We previously demonstrated that MYCN and PRMT5 proteins interact and PRMT5 knockdown led to apoptosis of MYCN amplified (MNA) neuroblastoma. Here we evaluate PRMT5 inhibitors GSK3203591/GSK3326593 as targeted therapeutics for MNA neuroblastoma and show MYCN-dependent growth inhibition and apoptosis. RNAseq revealed dysregulated MYCN transcriptional programmes and altered mRNA splicing, converging on key regulatory pathways such as DNA damage response, epitranscriptomics and cellular metabolism. Metabolic tracing showed glutamine metabolism was impeded following GSK3203591 treatment, which disrupted the MLX/Mondo nutrient sensors via intron retention of MLX mRNA. Glutaminase (GLS) protein was decreased by GSK3203591 despite unchanged transcript levels, suggesting post-transcriptional regulation. We demonstrate the RNA methyltransferase METTL3 and cognate reader YTHDF3 proteins are lowered following splicing alterations; accordingly, we observed hypomethylation of GLS mRNA and decreased GLS following YTHDF3 knockdown. In vivo efficacy of GSK3326593 was confirmed by increased survival of *Th-MYCN* mice together with splicing events and protein decreases consistent with in vitro data. Our study supports the spliceosome as a key vulnerability of MNA neuroblastoma and rationalises PRMT5 inhibition as a targeted therapy.

**GRAPHICAL ABSTRACT:** 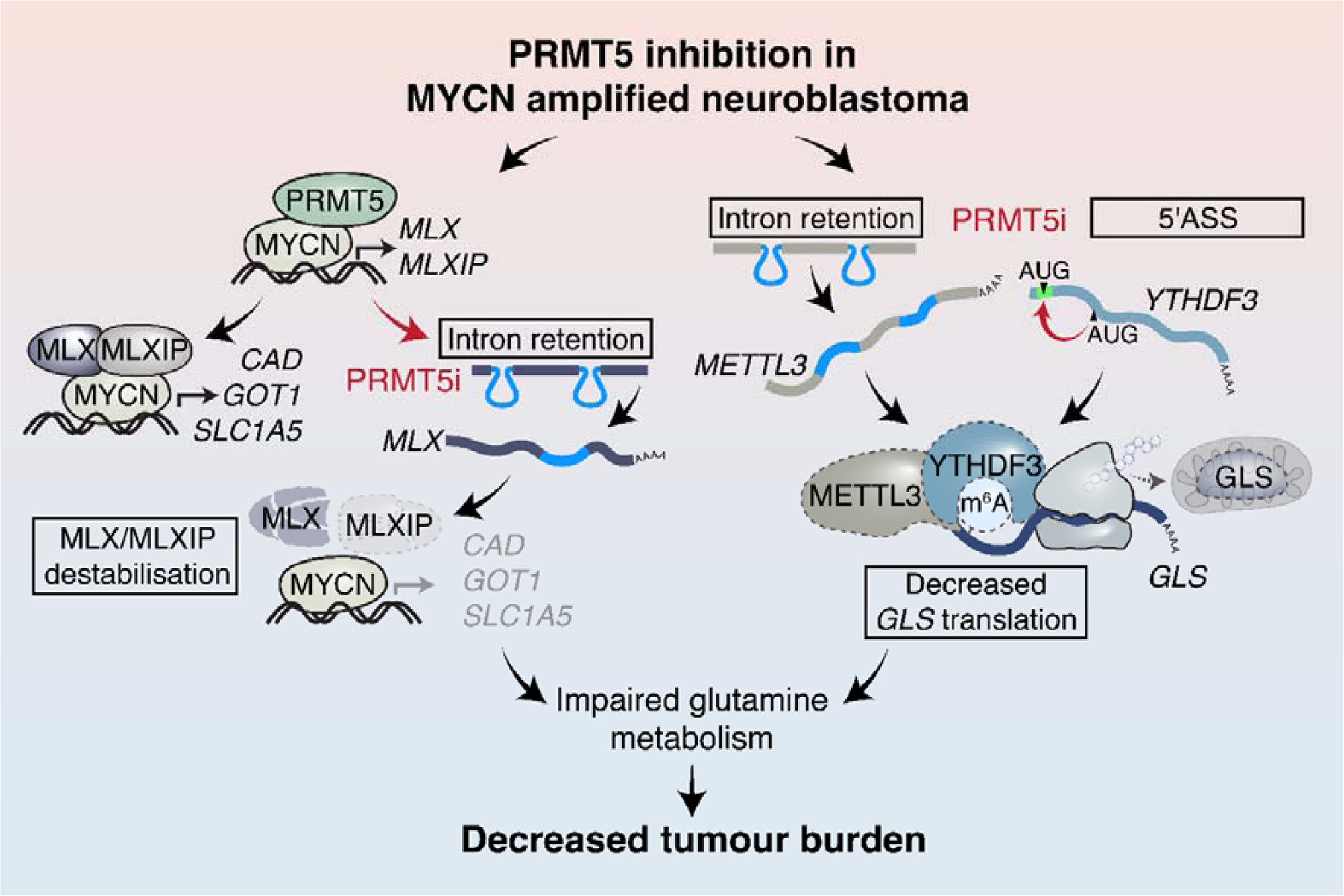

## INTRODUCTION

Neuroblastoma is one of the most common solid tumours of childhood, and approximately 50% of neuroblastoma patients have a high-risk clinical phenotype with very poor prognosis, specifically a long-term survival rate of less than 40% (1). The earliest defined driver of poor prognosis neuroblastoma is gene amplification of the *MYCN* proto-oncogene which encodes a transcription factor of the myc-family (2). *MYCN*-amplified neuroblastoma (MNA neuroblastoma) represents about half of poor prognosis neuroblastoma, with the remainder attributable to enhancer alterations leading to over-expression of *TERT* (3) or *MYC* (4).

Regrettably, these oncogenic drivers are not amenable to targeted therapies, underlining the crucial need for identifying druggable synthetic lethal or collateral vulnerabilities for efficacious therapies. Previous work in our laboratory demonstrated that MNA neuroblastoma cell survival was dependent on protein arginine methyltransferase 5 (PRMT5), with knockdown of PRMT5 resulting in MNA neuroblastoma cell line apoptosis (5). PRMT5 is one of two “type II” arginine methyltransferases, the other being PRMT9. Type II PRMTs catalyse symmetric arginine dimethylation (SDMA), in contrast to asymmetric arginine dimethylation (type I PRMTs) or monomethylarginine (type III PRMTs). PRMT5 and PRMT9 are not functionally redundant and exhibit very distinct substrate preferences (6). PRMT5 has pleiotropic functions and exerts oncogenicity by different mechanisms in many cancers (6,7), by functioning as a writer of symmetric dimethylation of the histone 3 at arginine 2 (H3R2me2s), arginine 8 (H3R8me2s) and histone 4 arginine 3 (H4R3me2s) marks associated with epigenetic silencing, exemplified by PRMT5 maintenance of breast cancer stem cells by epigenetically regulating *FOXP1* (8). Non-histone proteins are also substrates for PRMT5, with glycine-arginine rich (GAR) motifs being the preferred but not exclusive methylation sites and methylation by PRMT5 can alter the stability and activity of key transcription factors including p53 (9), E2F-1 (10,11), and MYCN in neuroblastoma (5). PRMT5 is a key regulator of mRNA constitutive and alternative splicing (AS) via arginine methylation of key spliceosomal components such as small nuclear ribonucleoproteins (U snRNPs) (12,13). Importantly, co-opting of spliceosomal regulation was shown to be essential for lymphomagenesis in an Eµ-myc driven mouse model (14), underlining the critical role of alternative splicing in cancer.

Given the evidence for PRMT5 representing a collateral vulnerability for the MYC-family proteins (5,14) together with the discovery of highly selective and potent PRMT5 small molecule inhibitors (15), we reasoned that pharmaceutical inhibition of PRMT5 may represent a promising targeted therapy for high-risk neuroblastoma. In this study, we report assessment of the substrate competitive PRMT5 inhibitors GSK3203591 and GSK3326593 *in vitro* and *in vivo* respectively. Our work demonstrates that PRMT5 is integral to survival and fitness programmes, including cellular metabolism, DNA repair and epitranscriptome modulation, via regulation of transcriptional and AS programmes in MNA neuroblastoma.

## RESULTS

### MYCN amplified neuroblastoma cell lines are preferentially sensitive to pharmaceutical inhibition of PRMT5

We sought to further develop our genetic interference data demonstrating PRMT5 requirement in MNA neuroblastoma (5) by evaluating the highly selective, first-in-class PRMT5 inhibitor GSK3203591 on a panel neuroblastoma cell lines. Initially we investigated growth inhibition using live-cell imaging of two MNA and two non-MNA neuroblastoma lines. Whereas the MNA lines IMR32 and SK-N-BE(2)-C (herein referred to as BE2C) demonstrated dose-dependent growth inhibition, the non-MNA lines SK-N-SH and SHIN did not (Figure 1A-B). We extended this analysis by determining IC_50_s for GSK3203591 in a panel of 14 neuroblastoma lines, together with three non-cancer lines RPE-1, NF-730 and NF-TERT. This verified that MNA-cell lines are >200 times more sensitive to the effects of PRMT5 inhibition compared to non-MNA lines, with mean IC_50_ values of 84 nM for MNA lines (range 33 nM – 144 nM) and 19.15 µM (range 8.3 µM – 29.4 µM) for non-MNA lines (Figure 1C). The non-cancer lines did not display IC_50_s in the concentration range 625 nM-80 µM used for non-MNA and non-cancer lines.

**Figure 1.**
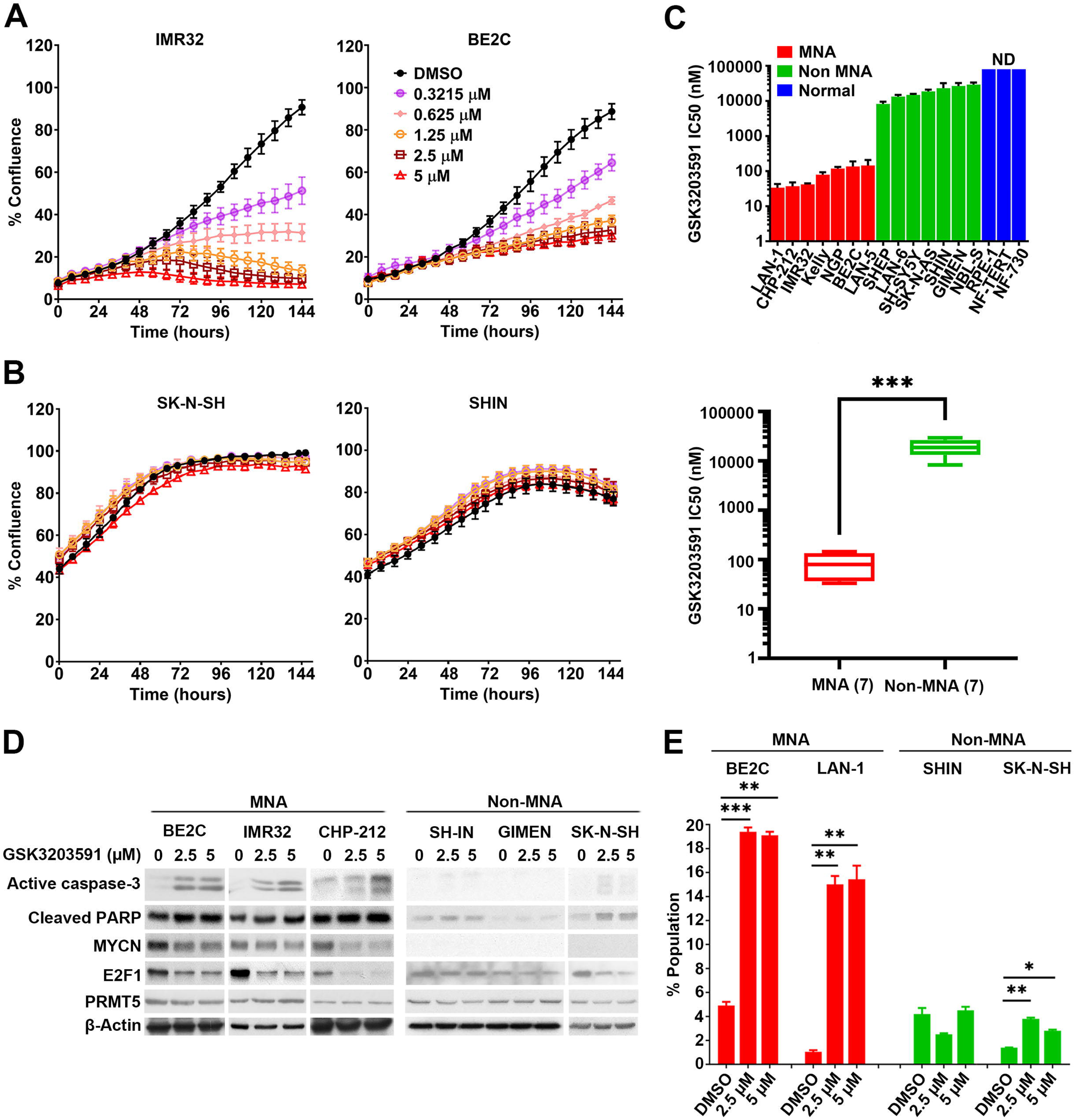
MYCN amplified cells are preferentially sensitive to PRMT5 inhibition. **[A]** Incucyte Zoom live cell imaging of two MYCN amplified (MNA) neuroblastoma lines demonstrating dose-dependent growth inhibition following GSK3203591 treatment (n=3, mean ± SD). **[B]** Imaging, as above, of two non-MNA neuroblastoma lines demonstrating no significant growth inhibition following GSK3203591 treatment (n=3, mean ± SD) **[C]**. *(Upper panel)* GSK3203591 IC_50_ values for 7 MNA, 7 non-MNA and 3 non-cancerous cell lines (n=3 ; mean ± SD, ND: not determined). *(Lower panel)* Box plot showing statistically significant mean IC_50_ ± SEM values of MNA vs non-MNA (***=p<0.0005, unpaired t-test). **[D]** Immunoblot of apoptotic markers, PRMT5, MYCN and E2F1 in cell extracts prepared from MNA (BE2C, IMR32, CHP-212) and non-MNA (SH-IN, GIMEN, SK-N-SH) neuroblastoma cell lines. β-actin was used as a control. One representative experiment from n=2 is shown. **[E]** Flow cytometry-based cell cycle analysis of two MNA (BE2C and LAN-1) and two non-MNA (SH-IN and SK-N-SH) cell lines, demonstrating increased sub-G1 apoptotic population (n=3, mean ± SD, *=p<0.05, **=p<0.005, ***=p<0.0005, student’s one-tailed t-test).

We next examined GSK3203591 effects on potential molecular targets and markers of apoptosis. Cleaved caspase-3 and cleaved PARP increased with GSK3203591 treatment in three MNA-lines, but barely changed in non-MNA cell lines (Figure 1D), although profound apoptosis was not observed with the treatment parameters. MYCN and E2F1 proteins were decreased, whereas PRMT5 levels were unchanged. All treatments were confirmed to have decreased symmetrical dimethyl arginine (SDMA) and not affected asymmetrical dimethyl arginine (ADMA) (Supplementary Figure 1). We further assessed apoptosis using cell cycle analyses to quantify the increased sub-G1 apoptotic population, confirming significant changes in the sub-G1 fraction on two MNA neuroblastoma lines, and minimal changes in the non-MNA lines, consistent with the immunoblots (Figure 1E). Together our analyses identify a *MYCN* amplification dependent sensitivity of neuroblastoma cell-lines to GSK3203591.

### Inducible expression of *MYCN* in isogenic SHEP-21N model sensitizes cells to GSK3203591

We next assessed the potential MYCN dependency of GSK3203591 sensitivity using isogenic SHEP-21N cells with tetracycline regulable MYCN (16). Proliferation assays revealed approximately 90-fold increased sensitivity with MYCN induced, i.e. an IC_50_ of 22.6±1.93 nM with MYCN induced and at 2.07±1.2 µM when MYCN is off (Figure 2A). Reflecting this, clonogenic assays demonstrated reduced colony formation in MYCN-on cells (Figure 2B). As the increased sensitivity to GSK3203591 could be attributable to the faster proliferation rate of MYCN-on cells, we also tested etoposide as a control. However, etoposide sensitivity did not segregate with MYCN expression status (Supplementary Figure 2A), further indicating that PRMT5 inhibition effects are highly specific to MYCN status. In line with this, apoptotic markers were more markedly increased in MYCN-on cells (Figure 2C). This trend was also apparent in cell cycle analyses revealing a significant increase in sub-G1 cells in MYCN-on cells, and not in MYCN-off cells. The cell cycle distribution analysis also indicated G1 arrest in MYCN-on cells after 48 hours treatment (Figure 2D).

**Figure 2.**
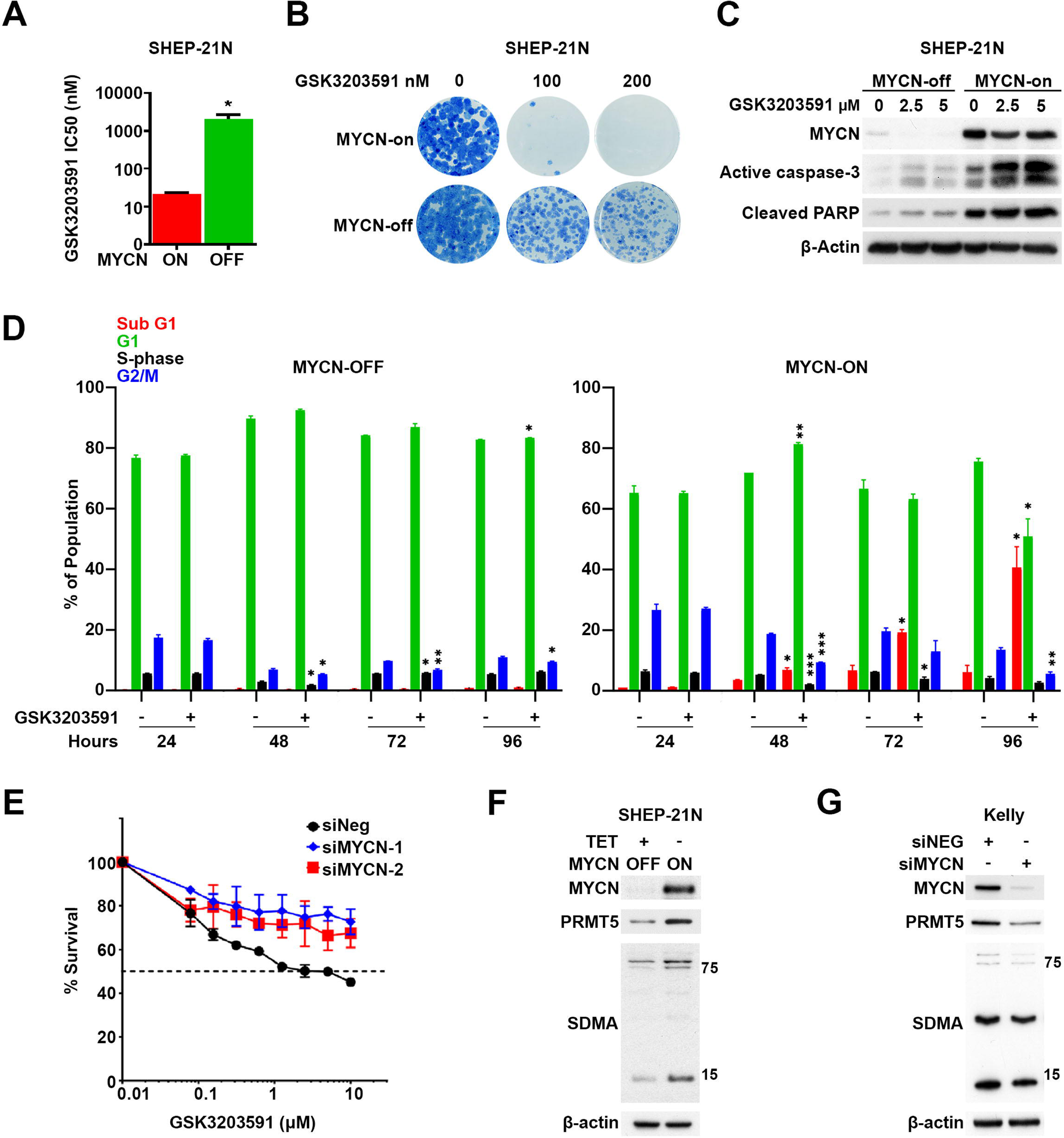
MYCN expression sensitizes neuroblastoma cells to PRMT5 inhibition. **[A]**. The isogenic neuroblastoma line SHEP-21N, with tetracycline regulable MYCN expression, shows increased GSK3203591 sensitivity when MYCN is switched on (n=3, mean ± SD, *=p<0.05, unpaired t-test). **[B]** Clonogenic assay with SHEP-21N cells demonstrating MYCN-dependency for GSK3203591-mediated growth inhibition. One representative experiment from 2 is shown. **[C]** Immunoblot analysis of cell extracts from SHEP-21N cells treated with GSK3203591 plus with (MYCN off) and without (MYCN on) tetracycline showing MYCN-dependent increases in apoptotic markers. β-actin was used as a control. One representative experiment from 2 is shown. **[D]** Flow cytometry cell cycle analysis by DNA content (propidium iodide) in SHEP-21N cells treated with 5 μM GSK3203591 or DMSO equivalent incubated with (MYCN off) and without (MYCN on) tetracycline for 24-96 hour time course (n=2, mean ± SD, *=p<0.05, **=p<0.005, ***=p<0.0005, unpaired t-test). **[E]** Cell survival assay (MTT) of BE2C cells transfected with two different siRNA’s targeting MYCN or siNEG and incubated with increasing concentrations of GSK3203591 for 96 hours (n=3, mean ± SD). **[F]** Immunoblot analysis of cell extracts from SHEP-21N cells incubated with (MYCN off) and without (MYCN on) tetracycline showing MYCN-dependent increases in PRMT5 and SDMA. β-actin was used as a control. One representative experiment from 3 is shown. **[G]** Immunoblot of cell extracts from Kelly cells transfected with siRNA targeting MYCN showing MYCN dependent PRMT5 and SDMA expression. β-actin was used as a control. One representative experiment from 2 is shown.

To further verify the MYCN-dependent sensitivity, we knocked down endogenous MYCN in BE2C cells and concurrently treated with GSK3203591 (Figure 2E). MYCN knockdowns increased resistance to GSK3203591, suggesting MYCN expression is essential for PRMT5 inhibition effects. As a control we also tested doxorubicin, but doxorubicin potency was not affected by MYCN knockdown (Supplementary Figure 2B). PRMT5 has been shown to be a c-myc target (14) and MYCN has been shown to bind the PRMT5 promoter (17), prompting us to evaluate MYCN effects on PRMT5 protein and symmetrical dimethylarginine (SDMA) using the SHEP-21N system. Induction of MYCN protein led to an increase in both PRMT5 and the SDMA mark (Figure 2F), and conversely, knockdown of MYCN in Kelly cells led to a decrease in PRMT5 and SDMA (Figure 2G).

Taken together, these studies verify that expression of MYCN protein in neuroblastoma cells is a major determinant of GSK3203591 sensitivity, and further validate a MYCN-PRMT5 axis crucial to survival and proliferation of neuroblastoma as suggested previously using PRMT5 knockdowns (5).

### PRMT5 inhibition leads to widespread gene expression changes, including MYCN-regulated genesets

To understand how GSK3203591 exerts growth inhibitory effects on MNA cell lines, we next sought to examine how PRMT5 inhibition may alter gene expression programmes in MNA lines. IMR32, Kelly and BE2C were treated with GSK3203591 and transcriptomes analyzed by RNA sequencing. We found 315 differentially expressed genes (DEGs, adjusted p<0.05, +/− 30% change) common to all 3 cell lines (Supplementary Figure 3A), and focused further investigation to this overlap set to find mechanistic determinants underscored by *MYCN* amplification and over-expression (Figure 3A). This set included 138 genes upregulated after GSK3203591 treatment and 177 genes downregulated genes (Supplementary File 1). A panel of 29 DEGs was validated by qRT-PCR in all 3 cell lines (Supplementary Figure 3B). Interestingly, scanning of expression levels and correlation with prognosis using the R2: Genomics Analysis and Visualization Platform (http://r2.amc.nl) suggested that an upregulated DEGs signature (DEG UP) did not significantly correlate with overall survival probability. Strikingly, the DEG DOWN signature showed a dramatic correlation with poor survival (p=9.7×10^−25^) (Figure 3B). Thus, high expression of the majority of genes from the DEG-DOWN signature associate with very poor prognosis in primary neuroblastoma, and PRMT5 inhibition lowers their levels in MNA neuroblastoma. These signatures suggest that PRMT5-mediated epigenetic silencing is unlikely to be the major determinant of MNA neuroblastoma sensitivity to GSK3203591. Conversely poor prognosis genes, potentially highly expressed because of MYCN-mediated transactivation, are mediators of increased survival and fitness of MNA neuroblastoma cells. The expression of these genes and their pathways are compromised by PRMT5 inhibition.

**Figure 3.**
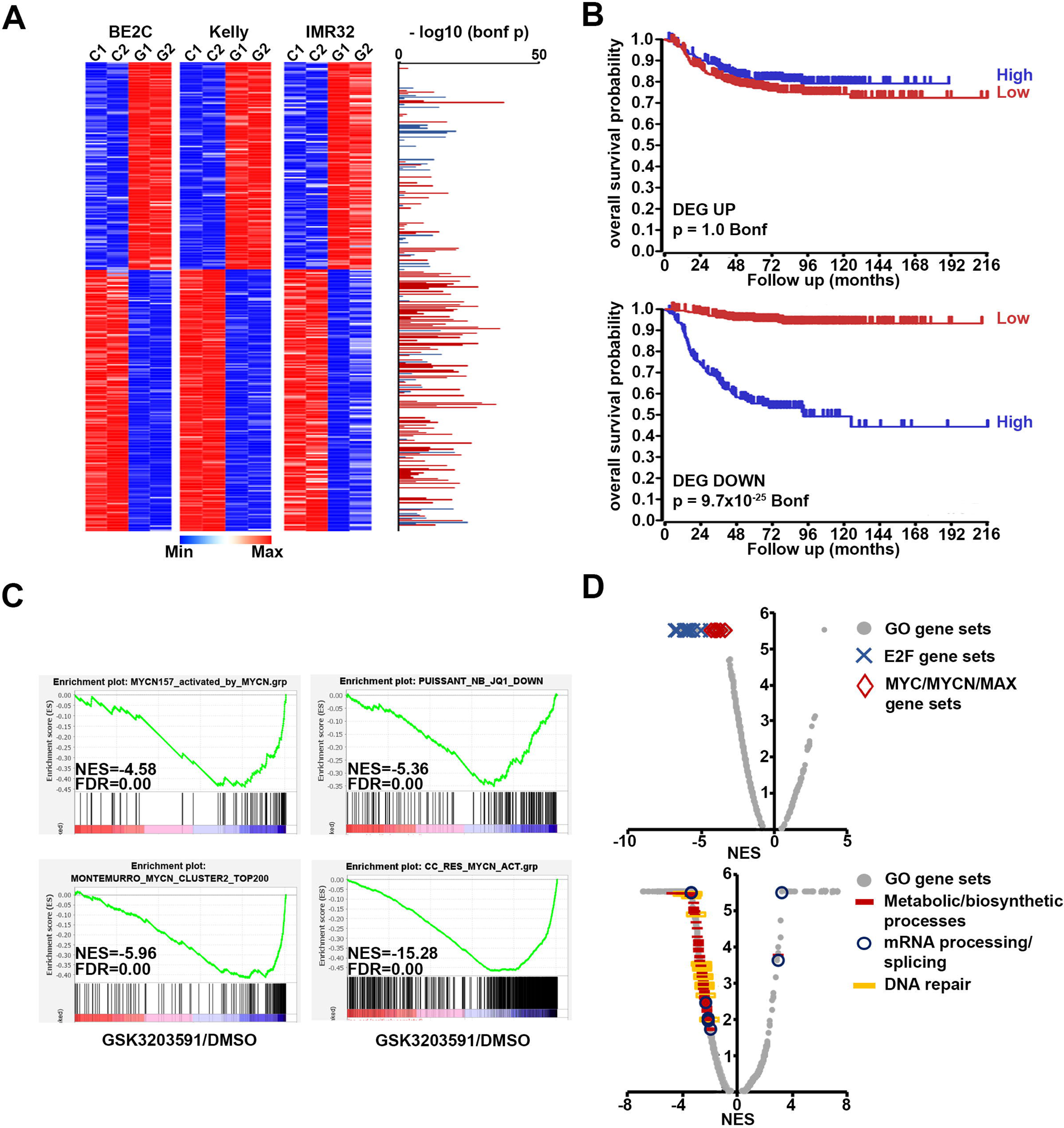
RNA sequencing of GSK3203591-treated MNA neuroblastoma lines reveals down-regulation of MYCN-activated genes. **[A]** Heatmaps of 315 differentially expressed genes (DEG, adjusted p<0.05, +/− 30% change) shared by BE2C, Kelly and IMR32 cell-lines following GSK3203591 treatment (C, control(DMSO); G, GSK3203591). Associations with prognosis for all genes is indicated alongside (right), bars indicate high expression is associated with poor prognosis (red) or good prognosis (blue). Bonferroni-corrected p-values were calculated on the R2 Genomics Analysis and Visualization Platform (http://r2.amc.nl) using the SEQC dataset containing gene expression data from 498 neuroblastoma patients. RNAseq was performed using n=2 biological replicates. **[B]** Kaplan Meier plots of gene signatures (metagenes) from genes upregulated following PRMT5 inhibition (DEG UP, top) or down-regulated (DEG DOWN, bottom) against overall survival. Note that high expression of DEG DOWN genes is strongly associated with poor survival. **[C]** GSEA analysis of GSK3203591-treated IMR32 cells demonstrating strong inhibition of MYCN-dependent gene sets. **[D]** Global summary of GSEA plots showing repression of MYC/MYCN and E2F gene sets (top) and other frequently altered gene ontology GSEAs affected by PRMT5 inhibition (bottom). NES, normalised enrichment score; FDR, false discovery rate.

We next conducted gene set enrichment analysis (GSEAs) to identify key pathways altered following PRMT5 inhibition. MYCN-activated genesets were strongly repressed in IMR32 cells (Figure 3C), as well as in Kelly and BE2C cell lines (Supplementary Figure 3C). Volcano plots of GSEAs indicated frequent decreases in MYC/MYCN/MAX and E2F genesets, the latter likely reflecting the regulation of E2F transcription factors by MYCN (18–20). Other notable changes (predominantly decreases) were observed in metabolic/biosynthetic process and mRNA processing, splicing and DNA repair (Figure 3D and Supplementary Figure 3D). Together, our RNAseq analyses show that GSK3203591 treatment suppresses fitness, survival and proliferation pathways, in particular MYCN-activated genes.

### PRMT5 inhibition leads to multiple alternative splicing events in MNA neuroblastoma

As PRMT5 is a key regulator of alternative splicing, further analysis of RNAseq data was conducted to identify differentially spliced genes (DSGs). Altogether 439 genes had at least 1 alternative splicing event shared between the 3 cell lines. These comprised of 32 genes with 3’-alternative splice sites (3’-ASS), 108 with 5’-alternative splice sites (5’-ASS), 101 with alternative cassette exons (CE), 62 with composite events (CompE) and 182 genes with ≥1 intron retention (IR) (Figure 4A, Supplementary File 1). We compared the potential association of DEGs and DSGs with neuroblastoma prognosis, specifically with the MYCN-157 gene signature that was found to have greater prognostic power than *MYCN* amplification status (21). As shown in Figure 4B, there are highly significant overlaps between DEGs, DSGs, PRMT5 expression versus the MYCN-157 signature, indicating that the genes which showed altered expression following PRMT5 inhibition, together with PRMT5, had significantly higher expression in MNA neuroblastoma and were co-expressed with the MYCN157 prognostic signature in the SEQC neuroblastoma dataset of 498 tumours. This shows that the PRMT5 regulated DEGs and DSGs strongly overlap with genes linked to poor prognosis. Reactome analysis highlighted RNA splicing/processing, cellular metabolism, and DNA repair as over-represented categories amongst downregulated DEGs and DSGs (Figure 4C). We first validated RNA splicing/processing hits identified amongst our DSGs; examples identified in our data include skipping of exon 8 in *HNRNPA1*, 5’-ASS of *HNRNPC*, and intron retention of *SRPK1* and *ZRANB2.* To our knowledge, these AS events have not been reported as being regulated by PRMT5 before (Figure 4D). Protein level changes of hnRNPA1 and SRPK1 following GSK3203591 treatment were also confirmed in MNA neuroblastoma cell-lines (Figure 4E). Interestingly, hnRNPA1 was previously demonstrated to be a MYCN target gene mediating AS in neuroblastoma (22), and a comparison of our DSGs with DSGs reported in RNAseq of hnRNPA1/PTBP1 knockdowns (22) demonstrated a significant overlap (Figure 4F). Several other genes encoding splicing and RNA metabolism regulators, including *PAPOLA, HNRNPD*, *HNRNPH1* and *PRPF3,* were also validated at the RNA level (Supplementary Figure 4A).

**Figure 4.**
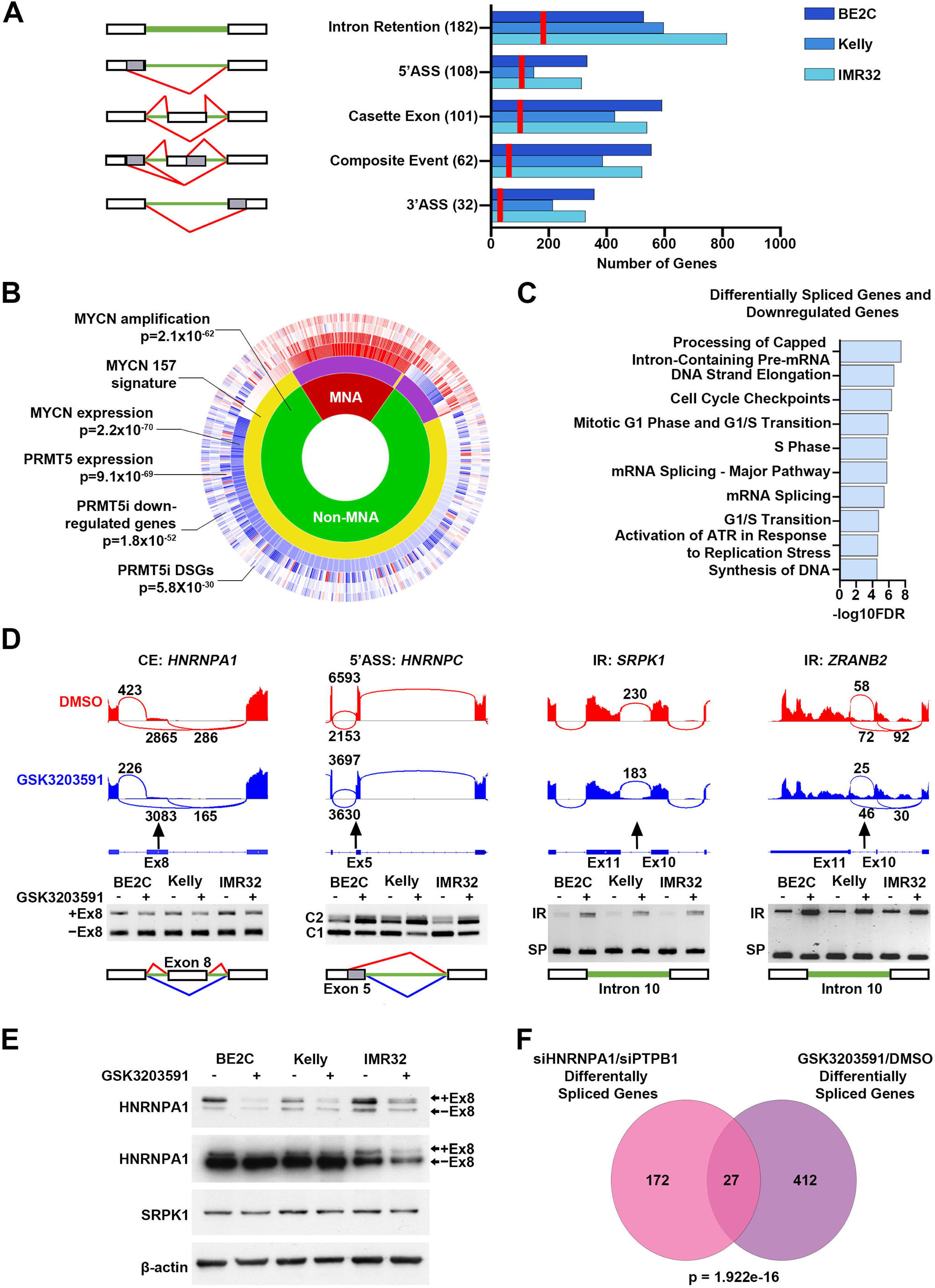
PRMT5 inhibition leads to widespread alternative splicing. **[A]** Numerical summary of alternative splicing events occurring in genes following PRMT5 inhibition in 3 MNA cell lines (BE2C, Kelly, IMR32) (right). Red lines on bar graph indicate the number of shared genes with alternative splicing events in the 3 cell lines, exact number displayed in brackets. Schematic (left) shows introns (green), exons (white) and splicing (red) (3’ASS, 3’alternative splice site; 5’ASS, 5’alternative splice site). **[B]** Sunburst plot generated in R2 (http://r2.amc.nl) showing, from inner to outer, correlation of *MYCN* amplification status (MNA/non-MNA), the MYCN-157 prognostic signature, *MYCN* transcript levels, *PRMT5* transcript levels, differentially expressed downregulated genes following PRMT5 inhibition (PRMT5i), and differentially spliced genes (DSGs) following PRMT5i. Overlap probabilities are shown relative to the MYCN-157 signature. **[C]** Top 10 significantly enriched Reactome pathways in a combined list of differentially spliced genes and downregulated differentially expressed genes. **[D]** Representative sashimi plots (top) and end-point PCR validation (bottom) of alternative splicing events occurring in genes that function in RNA splicing in BE2C, Kelly and IMR32 cells after GSK3203591 (IR, intron retention; SP, spliced product; CE, cassette exon; 5’ASS, 5’ alternative splice site; lines indicate splicing before (red) and after (blue) GSK3203591). **[E]** Immunoblot of cell extracts from BE2C, Kelly, IMR32 cells treated with GSK3203591 or DMSO equivalent (96h) showing decreased protein expression of hnRNPA1 (hnRNPA1B custom antibody targeting the + exon 8 isoform (top) and commercial hnRNPA1 antibody targeting the – exon 8 isoform (middle)) and SRPK1 (bottom). β-actin was used as a control. One representative of 3 experiments shown. **[F]** Venn diagram of differentially spliced genes in neuroblastoma cells transfected with siRNA targeting HNRNPA1/PTPB1 and differentially spliced genes shared by the 3 neuroblastoma cell lines following GSK3203591 treatment in this study (exact p value displayed).

Two recent studies demonstrated that targeting of the splicing regulator RBM39 using indisulam represents a novel therapeutic option for neuroblastoma (23,24). We therefore examined the extent to which our DSGs overlapped with DSGs following RBM39 knockdown in BE2C (23). Whilst a strikingly significant overlap was apparent between the two genesets, we observed no GSK3203591 effects on RBM39 protein levels (Supplementary Figure 4B-C).

PRMT5 inhibition has previously been reported to regulate the DNA damage response via alternative splicing of *KAT5* exon 5 (25). We also observed the AS of *KAT5* in our data (Supplementary Figure 4D) and in addition, we identified novel intron retention events in DNA repair genes *TIMELESS*, *PAXX* and *TOP2A*, as well as cassette exon splicing in *DONSON* (Figure 5A). We assessed the effect of PRMT5 inhibition on DNA damage repair in BE2C and Kelly cell lines by inducing double stranded DNA breaks by gamma irradiation after 48 hours GSK3203591 treatment. Increased DNA damage foci were observed in GSK3203591 treated BE2C and Kelly cells 1-hour post irradiation and DNA damage foci persisted 24-hours post irradiation in GSK3203591 treated BE2C cells (Figure 5B-C). These results show GSK3203591 impairs the DNA damage response and repair of double stranded DNA breaks in neuroblastoma.

**Figure 5.**
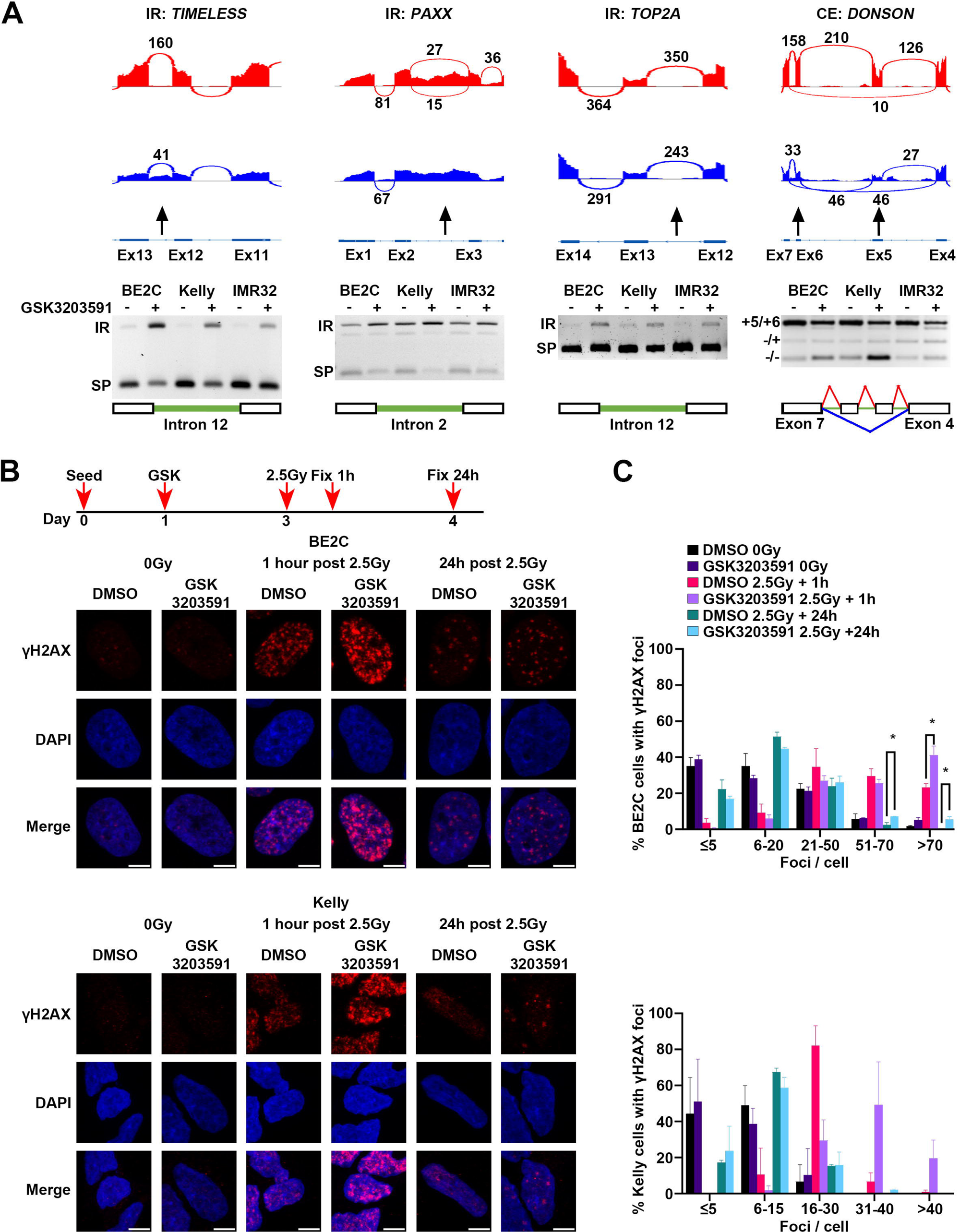
GSK3203591 induced alternative splicing of DNA repair factors. **[A]** Representative sashimi plots (top) and end-point PCR validation (bottom) of alternative splicing events occurring in genes that function in DNA damage response and repair (*TIMELESS, PAXX, TOP2A, DONSON*) in BE2C, Kelly and IMR32 cells treated with GSK3203591 or DMSO equivalent (96h) (IR, intron retention; SP, spliced product; CE, cassette exon; lines indicate splicing before (red) and after (blue) GSK3203591). One representative PCR experiment of 3 shown. **[B]** Representative confocal microscopy images (Z-projections) of γH2AX (red) in BE2C (top) and Kelly (bottom) cells treated with GSK3203591 and irradiated (nuclear counterstain Hoechst, blue; 63x magnification, scale bar 5 μM, representative images of three (BE2C) and two (Kelly) independent experiments). **[C]** Quantification of γH2AX foci per nucleus represented in B for BE2C (top, ≥125 cells counted from three independent experiments) and Kelly (bottom, ≥91 cells counted from two independent experiments) (mean ± SEM, unpaired t-test, *=p<0.05).

Together our AS analysis strongly supports deregulation of splicing programmes by PRMT5 as an important factor in neuroblastoma growth and survival.

### PRMT5 inhibition impacts neuroblastoma cellular metabolism

Numerous genes involved in metabolic pathways such as glycolysis and glutamine metabolism were apparent amongst the shared DEGs and DSGs, a selection of which are highlighted in Figure 6A. Notable DSGs included *PKM*, which has already been shown to undergo hnRNPA1-dependent alternative splicing in neuroblastoma (22) and *MLX*, encoding a MYCN transcriptional co-activator regulating glutamine metabolism genes in neuroblastoma (26). Switching between *PKM1* and *PKM2* isoforms involves inclusion of exon 9 (*PKM1*) or exon 10 (*PKM2*) and is associated with the Warburg effect in cancer cells (27). We confirmed AS of both *PKM* and *MLX*, together with the expected protein changes in all three MNA cell lines following GSK3203591 treatment (Figure 6B-D). We also identified decreases in Mondo/MLX interacting protein (*MLXIP),* previously shown to have interdependent expression with MLX (26), supporting the hypothesis that glucose and/or glutamine metabolism might be impaired following PRMT5 inhibition.

**Figure 6.**
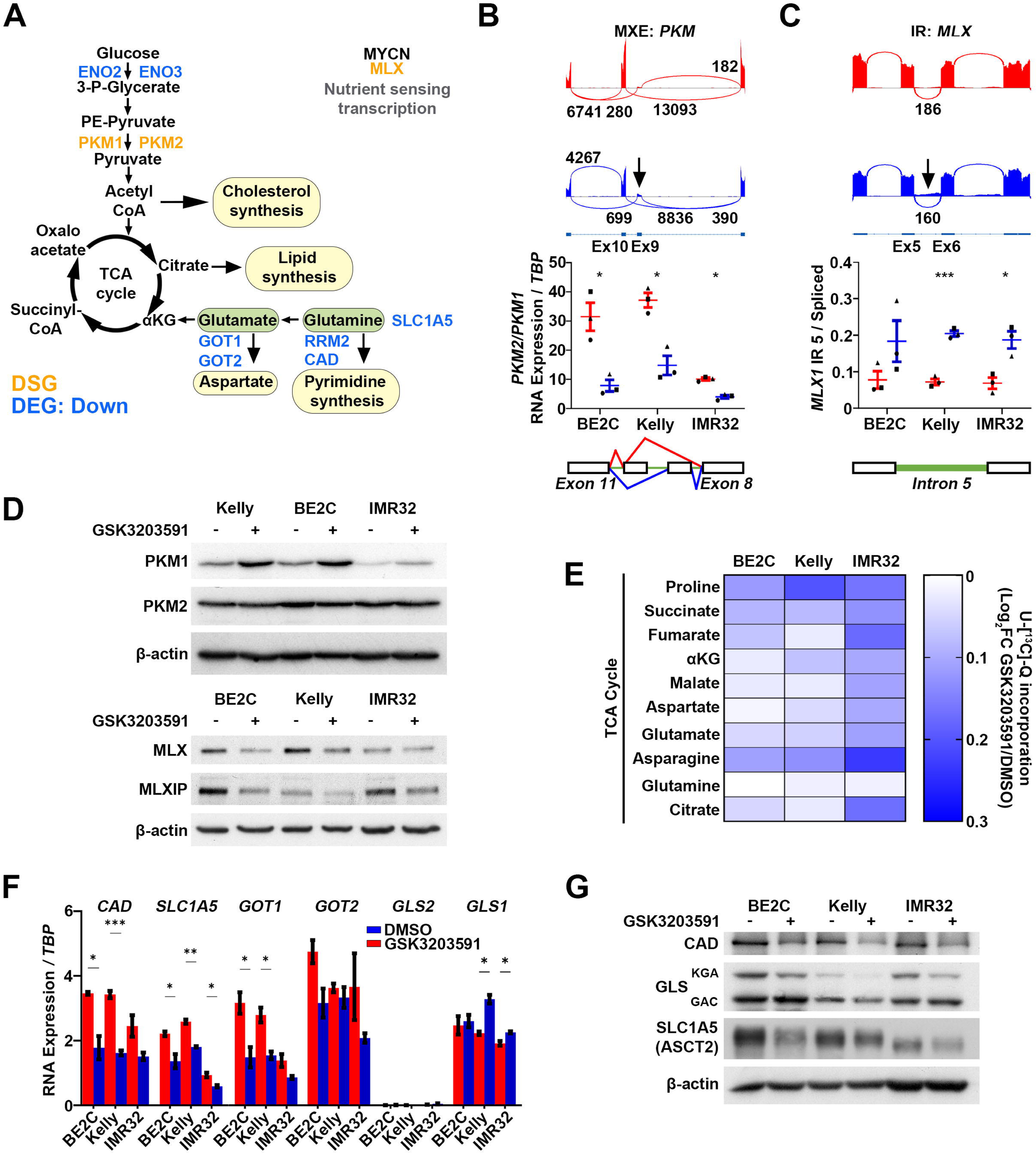
GSK3203591 induced metabolic changes. **[A]** Schematic diagram of glucose and glutamine metabolism highlighting differentially spliced genes (orange, DSG) and differentially expressed downregulated genes (blue, DEG Down) after GSK3203591 treatment. **[B]** Representative sashimi plot (top) and RT-qPCR validation (bottom) of PKM isoform expression (RT-qPCR data are relative to *TBP*, n=3 biological replicates, mean ± SEM, unpaired t-test, *=p<0.05) (MXE, mutually exclusive exon). **[C]** Representative sashimi plot (top) and RT-qPCR validation (bottom) of intron retention of intron 5 in the *MLX* transcript after GSK3203591 treatment (RT-qPCR data are relative to *TBP*, n=3 biological replicates, mean ± SEM, unpaired t-test, *=p<0.05, ***=p<0.0005, IR = intron retention) **[D]** Immunoblot of cell extracts from MNA cells treated with GSK3203591 showing PKM isoform switching from PKM2 to PKM1 (top) and downregulation of MLX and MLXIP expression (bottom). β-actin was used as a control. One representative experiment of 3 shown. **[E]** Heatmap summarizing glutamine incorporation into TCA cycle intermediates after PRMT5 inhibition relative to control in three cell lines (BE2C, IMR and Kelly), data derived from stable isotope labelling experiments (data represented as the average of 3 independent experiments). **[F]** RT-qPCR showing decreased expression of glutamine metabolism genes relative to *TBP* in MNA cell lines after GSK3203591 (n=3 biological replicates, mean ± SEM, unpaired t-test *=p<0.05, **=p<0.005, ***=p<0.0005). **[G]** Immunoblot showing decreased expression of glutamine metabolism proteins in MNA cell lines treated with GSK3203591. β-actin was used as a control. One representative of 3 experiments shown.

In order to test cellular utilization of glucose and glutamine following PRMT5 inhibition, we conducted stable isotope labelling using either uniformly labelled ^13^C_6_-glucose (U-[^13^C_6_]-Glc) or glutamine (U-[^13^C_5_]-Q). We observed decreases in glutamine-derived ^13^C incorporation in all three MNA neuroblastoma cell lines (Figure 6E and Supplementary Figure 5A and B), but no consistent changes in glucose incorporation into TCA intermediates following GSK3203591 treatment (Supplementary Figure 6A-C). We therefore validated genes encoding glutamine metabolism proteins in our treatments, and found downregulation of the glutamine transporter, *SLC1A5* and downstream glutamine metabolism genes *GOT1* and *CAD*. These genes have previously been linked with MYCN-associated glutamine addiction in neuroblastoma (28–30). Transcript levels of *GLS*, encoding glutaminase, the main pathway ‘gatekeeper’ enzyme, were either unchanged or slightly increased, and *GLS2* levels were very low in all three lines (Figure 6F). Immunoblotting confirmed that CAD and SLC1A5 protein levels were decreased after GSK3203591 treatment. Intriguingly, we found that GLS levels, specifically the GLS_KGA_ isoform, were decreased following PRMT5 inhibition (Figure 6G). GLS encodes two isoforms of glutaminase, kidney type glutaminase (KGA, 65 kDa) and glutaminase C (GAC, 58 kDa). However, GLS was not included in our shared DEGs or DSGs, suggesting that PRMT5 may regulate GLS via post-transcriptional mechanisms.

Overall, our metabolic studies strongly support the hypothesis that the PRMT5 dependency of MNA neuroblastoma is at least in part due to maintaining cancer cell fitness through glutamine addiction.

### PRMT5 inhibition leads to alterations in the epitranscriptome

The post-transcriptional regulation implied by our GLS analyses may occur via several mechanisms, including miRNA-mediated control or altered translational efficiency. Previous studies have shown c-myc regulates *AOPEP* which houses the GLS regulator miR-23 (31), however AOPEP was not in our DEGs. PRMT5 has already been shown to be involved in translational regulation via methylation of the hnRNPA1 protein, leading to enhanced hnRNPA1 binding to select internal ribosome entry sites (IRESs) and increasing translation (32). Another possible mechanism suggested by our DSG geneset was diminution of METTL3 as a result of increased intron retention following GSK3203591 treatment (Figure 7A). METTL3 is the key writer of m6A methylation on mRNA, which in turn lead to increased translation of mRNAs in cancer cells (33). We also identified an alternative 5’ splice site in the m6A reader YTHDF3 following GSK3203591 treatment (Figure 7A). We confirmed intron retention of introns 8 and 9 of *METTL3* following PRMT5 inhibition (Figure 7B) and validated YTHDF3 5’ splice site switching by end point PCR (Figure 7C). Genome database analysis of *YTHDF3* suggests that the alternative 5’ splicing results in switching from an open reading frame commencing in exon 3 to the canonical YTHDF3 using the exon 1 AUG codon. Analysis of universal RNA, fetal kidney, fetal adrenal and neural crest cell lines confirmed the generality of the YTHDF3 splice-isoforms (Supplementary Figure 7A). Alternative splicing events were accompanied by decreased METTL3 and YTHDF3 protein (Figure 7D); whereas METTL3 reduction is predicted from introducing premature termination codons, the mechanisms by which YTHDF3 translational codon switching reduce protein remain to be investigated.

**Figure 7.**
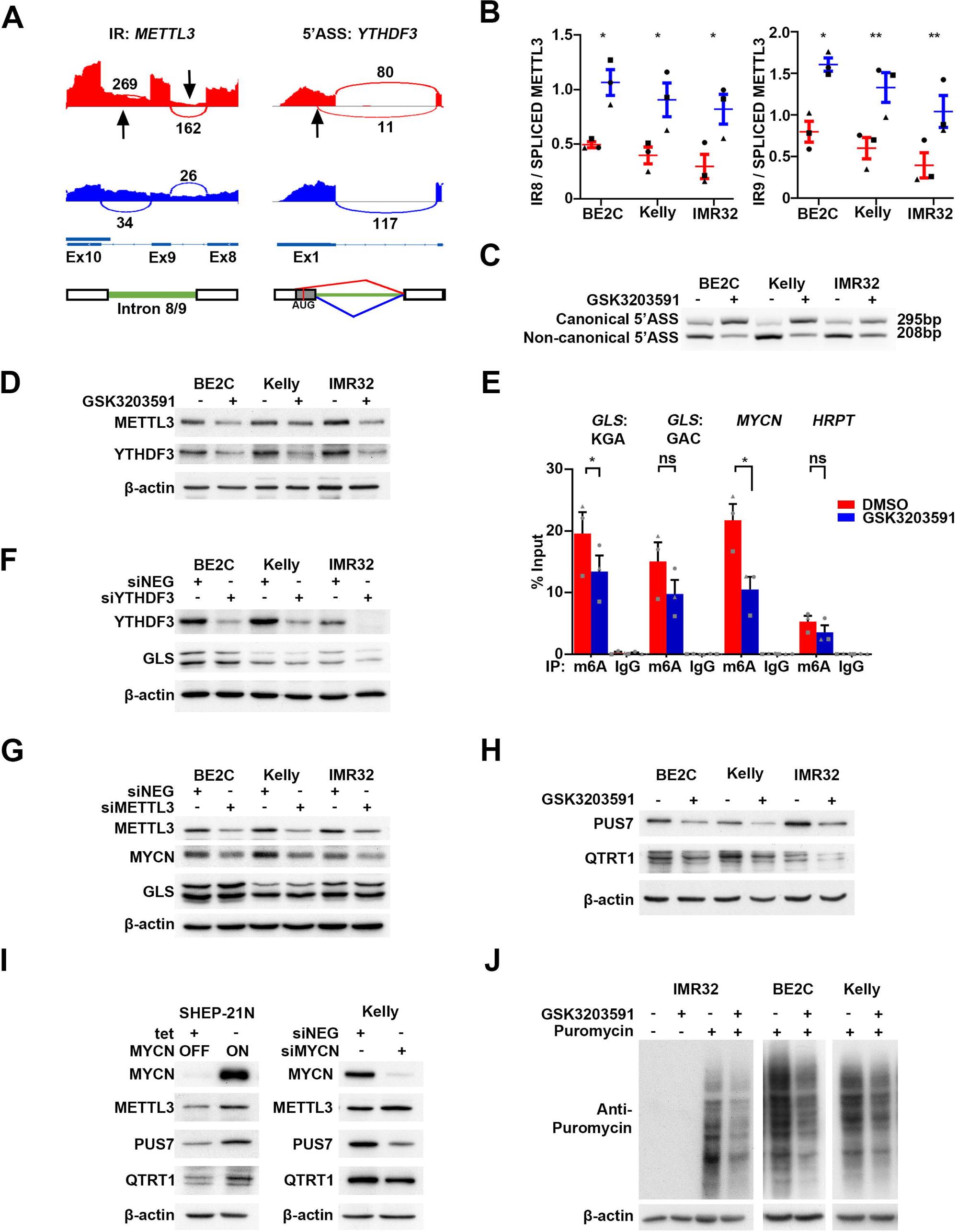
PRMT5 inhibition alters epitranscriptome regulators and decreases efficiency of translation. **[A]** Representative sashimi plots of METTL3 intron retention of intron 8 and 9 (left) and YTHDF3 alternative 5’ splice site in exon 1 (right) after GSK3202591. **[B]** RT-qPCR validation of increased intron retention of introns 8 and 9 in the *METTL3* transcript (n=3 biological replicates, paired t-test, *=p<0.05, **=p<0.005). **[C]** End point PCR validation of alternative 5’ splice site in exon 1 of YTHDF3 from the non-canonical splice site (DMSO) to the canonical splice site (GSK3203591). One representative experiment of 3 is shown. **[D]** Immunoblot showing decreased expression of METTL3 and YTHDF3 in BE2C, Kelly and IMR32 cells after GSK3203591. β-actin was used as a control. One representative experiment of 3 is shown. **[E]** Methylated RNA immunoprecipitation (MeRIP) performed using anti-m6A antibody demonstrates decreased m6A on *GLS* and *MYCN* transcripts detected by RT-qPCR in BE2C cells. HRPT was used as a control. (n=3 biological replicates, mean ± SEM, paired t-test, *=p<0.05). **[F]** Immunoblot showing decreased expression of GLS following YTHDF3 knockdown. β-actin was used as a control. One representative experiment of ≥2 shown. **[G]** Immunoblot showing decreased expression of MYCN following METTL3 knockdown. One representative experiment of ≥2 shown. **[H]** Immunoblot showing decreased expression of RNA modifier proteins PUS7 and QTRT1 in BE2C, Kelly and IMR32 after GSK3203591. β-actin was used as a control. One representative experiment of 3 is shown. **[I]** Immunoblot of cell extracts of SHEP-21N incubated with (MYCN off) or without (MYCN on) tetracycline (left) and Kelly cells transfected with siRNA targeting MYCN (right) probing for RNA modifier proteins PUS7, QTRT1 and METTL3. β-actin was used as a control. One representative experiment of 3 shown. **[J]** SunSET assay showing decreased protein translation after GSK3203591. β-actin was used as a control. One representative experiment of 2 is shown.

We therefore assessed whether *GLS* mRNA was marked by m6A methylation, and whether this epitranscriptomic mark is altered by PRMT5 inhibition. As shown in Figure 7E, m6A-methylated mRNA immunoprecipitation (MeRIP) confirmed both methylation of GLS mRNA and its reduction after GSK3203591 treatment of BE2C cells. GLS^KGA^ mRNA had higher m6A levels and a more significant decrease than GLS^GAC^, consistent with decreases in GLS^KGA^ protein observed in Figure 6G. As GLS has been previously demonstrated to regulated by YTHDF1 (34) we knocked down YTHDF3 and discovered decreased GLS protein expression (Figure 7F). These data demonstrate GLS is regulated by m6A modification and that the YTHDF3 reader is required for GLS translation.

Methylation of *MYCN* mRNA by METTL3 has been demonstrated recently (35), and we also observed a significant decrease of m6A methylation of *MYCN* mRNA following PRMT5 inhibition (Figure 7E). We next sought to determine the effect of METTL3 knockdown on MYCN expression and identified decreased MYCN protein expression following knockdown (Figure 7G). Developing on this evidence for post-transcriptional modifications, we further examined the intriguing link between PRMT5-MYCN and the epitranscriptome. Surveying of our DEGs and DSGs revealed potential decreases in several genes encoding epitranscriptomic modifiers, including the DEG *PUS7*, as well as the intron retention DSGs *QTRT1*, *NOP2* and *NSUN2* (supplementary Figure 7B-C). Interestingly *PUS7* is already identified as a MYCN-activated gene within the poor prognosis MYCN-157 signature (21). Immunoblotting confirmed decreases of PUS7 (regulator of tRNA and mRNA pseudouridylation) and QTRT1 (catalyzes the tRNA base-exchange of guanine with queuine) proteins following PRMT5 inhibition (Figure 7H). Epitranscriptome modifier protein decreases following *MYCN* knockdown or increases accompanying *MYCN* induction in SHEP-21N cells were also confirmed for PUS7 and QTRT1 (Figure 7I), further tightening the functional association of PRMT5-MYCN in post-transcriptional regulation.

As PRMT5 inhibition was demonstrated to have marked effects on epitranscriptomic regulators, we next explored whether global translation was affected by GSK3203591 in MNA neuroblastoma cells. For this we used the surface sensing of translation (SUnSET) assay which relies on incorporation of puromycin, a structural analogue of aminoacyl tRNAs, into nascent polypeptide chains, followed by probing cellular proteins with anti-puromycin antibodies. GSK3203591 consistently led to decreased puromycin incorporation, consistent with a global inhibition of translation and reduced cellular metabolism by PRMT5 inhibition (Figure 7J).

Taken together, our *in vitro* data demonstrate that PRMT5 supports the fitness, survival, and proliferation of MNA neuroblastoma via diverse mechanisms including previously reported pathways such as mRNA splicing and DNA repair, but also novel routes such as MYCN-mediated transcriptional regulation, glutamine metabolism and epitranscriptome regulation.

### Inhibition of PRMT5 *in vivo* significantly increases survival in a murine neuroblastoma model

Following our *in vitro* analyses, we assessed the *in vivo* efficacy of PRMT5 inhibition. For this we used the *Th-MYCN* mouse model for neuroblastoma, and an analogue of GSK3203591, namely GSK3326593, which has better characteristics for *in vivo* work. The *Th-MYCN* GEMM was generated by targeting MYCN expression to the neural crest under the regulation of the tyrosine hydroxylase promoter. Importantly, these tumours faithfully recapitulate the pathologic and molecular features of the human disease and have been used extensively to evaluate novel therapeutic strategies aimed at treating the poor-outcome group of neuroblastoma patients (36). A significant improvement in survival rates was apparent in mice dosed with 100 mg/kg GSK3326593 twice a day (Log-rank (Mantel-Cox) test, p=0.0265) (Figure 8A). Parallel treatments for pharmacokinetic/pharmacodynamic studies were harvested following five days of GSK3326593 or vehicle treatment and used for molecular analyses. Immunoblotting with anti-SDMA antibodies showed clear decreases in SDMA, confirming the on-target effect of GSK3326593, together with a small but consistent decrease of Mycn protein. Apoptosis markers were not increased following Prmt5 inhibition, but γ-H2ax increased, indicating elevated DNA damage (Figure 8B). We confirmed several of our DSGs in mouse tumours, including *Hnrnpa1*, *Srpk1*, *Hnrnpc*, and *Mlx* (Figure 8C). We further demonstrated decreases of glutamine pathway enzymes Gls and Cad proteins, although *Gls* mRNA was unaltered (Figure 8D). We therefore assessed Mettl3 protein and *Mettl3* intron retention and observed trends of decreased Mettl3 protein and increased *Mettl3* intron retention following GSK3326593 treatment (Figure 8E), consistent with our findings in MNA cell-lines, where GLS expression is subject to epitranscriptomic post-transcriptional control.

**Figure 8.**
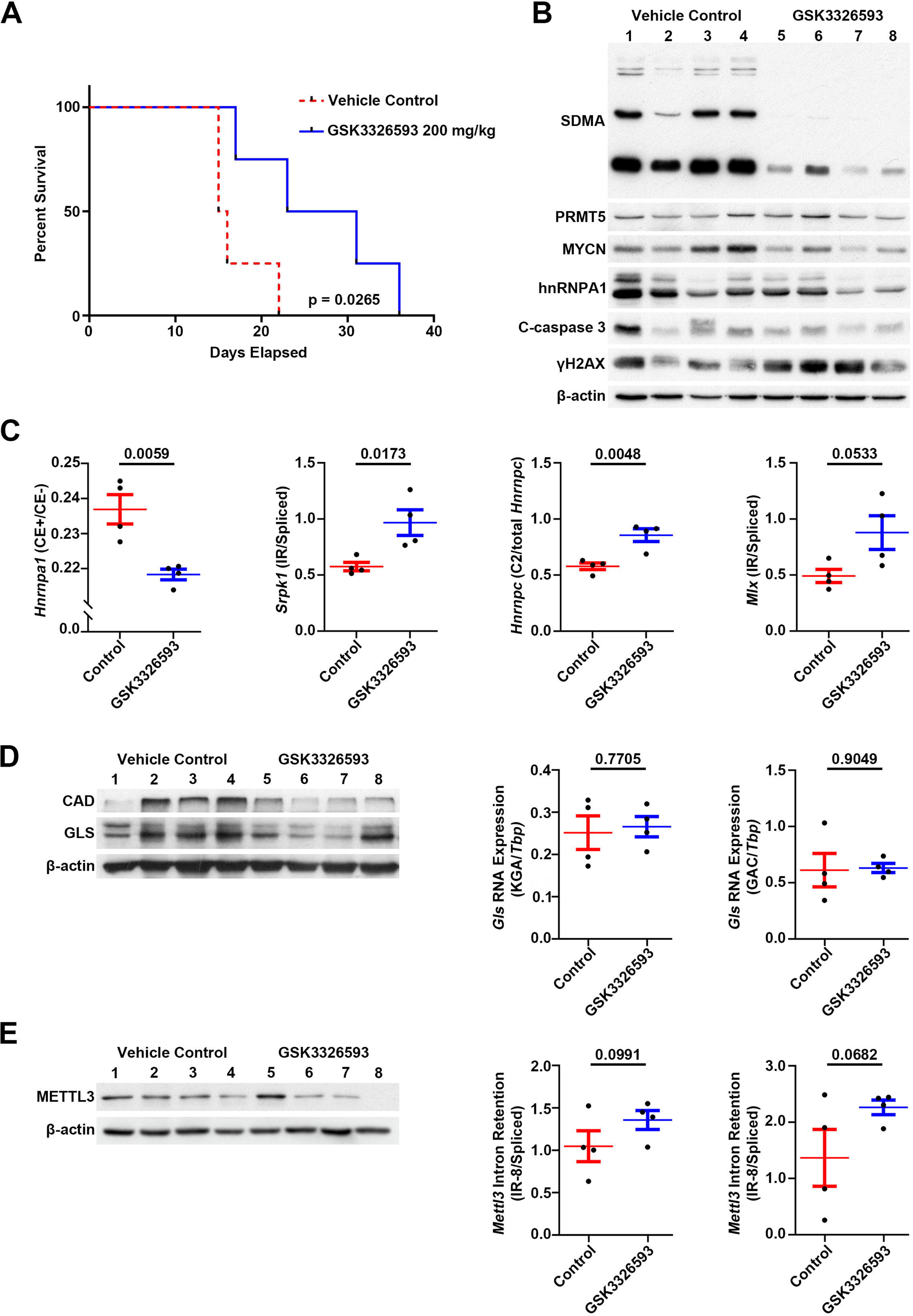
*In vivo* inhibition of PRMT5 in the *Th-MYCN* mouse neuroblastoma model significantly increases survival. **[A]** Kaplan-Meier showing survival of mice treated with GSK3326593 at 200 mg/kg (100 mg/kg twice a day) until maximum tumour burden was reached. Statistical significance for survival rates was determined using Log-rank (Mantel-Cox) test. Control cohort, n=4; treated cohort n=4. **[B]**. Immunoblot of protein extracts of mouse tumours showing inhibition of global SDMA in GSK3326593 treated tumours confirming target engagement (PRMT5) of GSK3326593. Immunoblots were probed with the indicated antibodies and β-actin was used as a control. **[C]** RT-qPCR, of a selection of alternative splicing events first identified *in vitro,* using RNA extracted from Th*-MYCN* mouse tumours after PRMT5 inhibition. Data relative to housekeeping gene *Tbp*. Control cohort, n=4; treated cohort n=4; unpaired t-test, exact p-value displayed. **[D]** Immunoblot of protein extracts of mouse tumours from control and GSK3326593 treated mice showing decreased GLS and CAD after GSK3326593 treatment (left), β-actin was used as a control. RT-qPCR (right) shows no significant change of *Gls* isoforms KGA/GAC mRNA, data relative to housekeeping gene *Tbp*. Control cohort, n=4; treated cohort n=4; unpaired t-test, exact p-value displayed. **[E]** Immunoblot (left) of protein extracts from mouse tumours showing decreased METTL3 protein after GSK3326593 treatment, β-actin was used as a control. RT-qPCR (right) of RNA extracted from mouse tumours demonstrating increased intron retention of intron 8 and 9 in the *Mettl3* transcript. Data relative to housekeeping gene *Tbp*. Control cohort, n=4; treated cohort, n=4; unpaired t-test, exact p-value displayed.

In summary, our *in vivo* data demonstrate that GSK3326593 is a highly selective inhibitor, and that inhibition of Prmt5 in Mycn-dependent mouse neuroblastoma compromises the fitness and proliferation of tumour cells, resulting in their increased survival.

## DISCUSSION

The importance of PRMTs in tumourigenesis is reflected in the increasing interest in developing targeted therapeutics for this family of enzymes, with several studies demonstrating that PRMTs are a synthetic or collateral vulnerability in cancer (37). The association of PRMTs with key oncogenes is best demonstrated in *Eµ-myc* driven lymphomagenesis in transgenic mice where c-myc requires Prmt5-mediated regulation of the spliceosome (14), and PRMT5 also acts with several other oncogenic pathways (38). In the case of solid tumours, our group previously demonstrated that neuroblastoma cells over-expressing the *MYCN* oncogene are highly susceptible to loss of PRMT5 (5), prompting this preclinical assessment of the selective PRMT5 inhibitors GSK3203591 and GSK3326593 in neuroblastoma. We show that MNA neuroblastoma is indeed sensitized to PRMT5 inhibition *in vitro* and *in vivo*, and identify further actionable pathways through our mechanistic analyses, for example epitranscriptomic and metabolic pathways.

Analysis of our neuroblastoma cell line panel demonstrated increased sensitivity of MNA neuroblastoma lines to GSK3203591, irrespective of their *TP53* mutational status. For example, CHP-212, IMR32 and NGP cell lines have wild-type *TP53*, whereas others such as LAN-1, Kelly and BE2C have mutant *TP53.* Activation of the p53 pathway was previously shown to increase sensitivity to GSK3203591 due to alternative splicing of *MDM4* transcripts induced by PRMT5 inhibition, leading to skipping of exon 6 which encodes the p53 interaction domain (39). Whilst we observed exon skipping of exon 6 of *MDM4* mRNA in neuroblastoma cell lines too, our GSK3203591 sensitivity spectrum suggests that p53 activation is not the major determinant of drug sensitivity in MNA neuroblastoma. Whilst PRMT5 levels were unchanged following GSK3203591 treatments, we observed decreases of MYCN and E2F1. PRMT5 was previously shown to methylate and stabilize E2F1 (10) so the observed decrease is likely attributable to altered MYCN regulation of *E2F1* which is an established MYC/MYCN target gene (18–20). Significantly more apoptosis was seen in MNA cell-lines, although overall cell death was modest even in sensitive cell-lines. The MYCN-dependency for GSK3203591 sensitivity is further strengthened by the SHEP-21N isogenic cell-line model with tetracycline-regulated MYCN where high MYCN led to decreased clonogenic survival, increased apoptosis and elevated PRMT5 and SDMA. PRMT5 has previously been shown to be a c-myc (14) and MYCN target (17), consistent with our protein level evidence. Together, our genetic interference (5) and pharmacological inhibition data strongly suggest that PRMT5 is synthetically lethal with MYCN in neuroblastoma. Apart from *TP53* status (discussed above), the most compelling PRMT5 collateral vulnerability reported to date is with loss of the *MTAP* (methylthioadenosine phosphorylase) gene leading to elevation of cellular methylthioadenosine, resulting in PRMT5 inhibition (40,41). However, *MTAP* lesions have not been shown in neuroblastoma – in fact *MTAP* has been reported to be transactivated by MYCN (21). It will therefore be important to evaluate the dependency of other MYCN-driven malignancies on PRMT5 to establish the generality of the MYCN-PRMT5 axis vulnerability.

Inhibition of PRMT5 led to a dramatic curtailment of *MYCN*-activated genes, including gene signatures strongly associated with poor prognosis in neuroblastoma (21,42), BRD4 inhibition of neuroblastoma with JQ-1 (43) and cell-cycle resolved MYCN-activated genes from RNA sequencing in cell-cycle-synchronized neuroblastoma cells (44). Whilst the number of genes whose transcripts increased after GSK3203591 treatment was similar to decreased transcripts, meta analysis with gene signatures strongly suggested that the major determinants for GSK3203591-mediated growth inhibition were the genes decreasing after PRMT5 inhibition. PRMT5 is known to exert epigenetic silencing through histone 3 arginine methylation (7), but our DEGs suggest that epigenetic derepression via PRMT5 inhibition is not a major route of drug action in MNA neuroblastoma. Global decreases of the PRMT5 histone modifications were also previously shown to be unaffected by GSK3203591, although specific loci may be epigenetically regulated (15). GSK3203591 treatment led to slight decreases of MYCN protein *in vitro* and *in vivo*, but not dramatically as we previously observed after PRMT5 knockdowns (5). This suggests that the PRMT5-MYCN protein interaction likely stabilizes MYCN independent of PRMT5 catalytic activity similar to the manner in which Aurora kinase A (45) and EZH2 (46) have been shown to stabilize MYCN protein. Activity of the MYC-family transcription factors can be influenced by many interactions including MAX, MXD1-4 and MNT (47) but these were not altered amongst our DEGs following GSK3203591 treatment. However, we did observe altered splicing of another MYCN transcriptional co-factor MLX (discussed below) which may explain, at least in part, the profound effect of PRMT5 inhibition on MYCN transcriptional programmes.

PRMT5 is known to be crucial in the assembly of spliceosomal complexes that regulate alternative splicing. Key substrates for PRMT5 include the Sm proteins, and restriction of their symmetrical dimethylation results in incorrect assembly into snRNPs. This results in the failure of complexes to recognize weak 5’-splice sites, inducing a variety of splicing defects, in particular intron retention (13,14). Consistent with this, GSK3203591 treatment resulted in numerous alternative splicing events shared between our three MNA cell-lines, approximately half of which were intron retentions. Similar predominance of intron retention following pharmaceutical inhibition of PRMT5 has been shown in glioblastoma and haematopoietic cells (48,49). There was a very high correlation between our DEGs and DSGs and the MYCN-157 signature. This signature was reported as a stronger prognostic indicator than MNA alone, and our data shows correlations where *MYCN* transcript levels were relatively low but MYCN protein levels were significant (21). Whilst high myc levels have also been shown to correlate with high-risk neuroblastoma (4), positive *MYC* correlation with our DEGs or DSGs was not observed.

The JUM bioinformatic pipeline yielded strongly validated alternative splicing hits encompassing many key regulatory pathways, including DNA repair and mRNA processing, as shown in glioblastoma (48). The multifunctional protein hnRNPA1, already known to be post-translationally modified by PRMT5, CARM1 and PRMT7 (32,50), was consistently shown to switch from an alternatively spliced (+ exon 8) isoform to a minus exon 8 form at RNA and protein level following GSK3203591 treatment. Whilst little is known about this isoform, it has been demonstrated to have much stronger binding to RNA (51) and also be expressed at elevated levels in chronic myelogenous leukemia compared to control cells (52). Given that hnRNPA1 is involved in many essential pathways including transcriptional and translational regulation, splicing and telomere maintenance (53), it will be of interest to examine the tumorigenic role of hnRNPA1(+ exon 8) in future. Alternative splicing of several other *HNRNP* genes was also apparent, suggested that PRMT5 regulates the intricately regulated assembly of hnRNP complexes that regulate splicing (54).

PRMT5 regulates DNA repair through multiple mechanisms such as direct methylation of RUVBL1 (55) and alternative splicing of *KAT5* (56). Our study reveals several other genes encoding DNA repair pathway proteins undergo splicing alterations that accompany enhanced DNA damage by GSK3203591, including *TIMELESS* (57) and *DONSON* (58). PRMT5 inhibition has recently been shown to enhance the sensitivity of breast and ovarian cancers to PARP inhibitors (59), and the convergence of alterations in DNA replication and repair genes observed in our data suggests that similar combinations could be effective against poor prognosis neuroblastoma.

Having shown that PRMT5 inhibition triggers widespread splicing changes, possibly through master nodes such as hnRNP proteins, we endeavoured to characterize key downstream events. As we mostly observed inhibition of cell growth rather than pronounced apoptosis, suggesting that PRMT5 may support the metabolic fitness of MNA neuroblastoma, we were intrigued by the consistent PKM2-PKM1 splicing switch observed after PRMT5 inhibition. PKM2 drives aerobic glycolysis in preference to oxidative phosphorylation (27) and the isoform switch has been shown to depend on hnRNPA1 (22,60). Despite strong mRNA and protein validation of the PKM isoform switching, metabolic tracing did not show any consistent effects on glucose metabolism. However, PKM2 also phosphorylates non-metabolic substrates, for example stabilizing Bcl-2 (61) and influencing transcription via phosphorylation of histone 3, already demonstrated in neuroblastoma (62), and perturbation of these pathways may contribute to GSK3203591-mediated growth inhibition. Numerous other DEGs and DSGs identified by our transcriptomic analyses encode metabolic pathway proteins. Given the MNA neuroblastoma-specific inhibition spectrum exhibited by GSK3203591 and the addiction of MNA neuroblastoma to glutamine (28,63), the convergence of DEGs and DSGs on genes in this pathway strongly supports the premise that PRMT5 contributes to the metabolic fitness of MNA neuroblastoma via augmenting glutamine metabolism. MYCN-dependent transcription of genes such as *SLC1A5*, *CAD* and *GOT1* has been previously reported although regulation of the main pathway ‘gatekeeper’ *GLS* has not. Although c-myc has been shown to regulate GLS expression via miR-23 (31), we note that miR-23 levels are not significantly different in a comparison of MNA and non-MNA neuroblastomas (64). Whilst we do not exclude this possibility, our studies demonstrate that GLS levels are indeed regulated by MYCN-PRMT5 through alternative splicing and epitranscriptomic pathways via the m6A reader YTHDF3. PRMT5 has recently been shown to control intron retention of METTL3 in blastic plasmacytoid dendritic cell neoplasm (65), similar to our observations in neuroblastoma. Interestingly, METTL3 knockdown alone did not cause GLS protein to decrease, possibly because METTL3 loss may cause increased 3’-UTR lengthening of GLS as previously reported (66) thus favouring KGA isoform expression. The combination of decreased *GLS* mRNA m6A-methylation and decreased YTHDF3 protein expression support translational inhibition of *GLS* mRNA by GSK3203591 contributing to altered glutamine metabolism. Whilst the read-outs of epitranscriptome marks such as m6A-methylation are very diverse (67), epitranscriptomic and mRNA translation control of GLS is supported by YTHDF1-dependent GLS regulation in colon cancer (34). Notably, our work suggests that PRMT5 is at the crux of epitranscriptome regulation as we also observed numerous other RNA writers and readers as targets for PRMT5-dependent splicing, including *QTRT1*, *NOP2* and *NSUN2*. The multiple levels of RNA regulation via these proteins, and of the hnRNPs, also known to interact with the epitranscriptomic machinery (68), suggests that the PRMT5-MYCN axis is also integral in proteostasis in neuroblastoma, which is supported by our global translation analyses. Protein translation is one of the highest energy consuming processes in the cell, and is therefore also likely to be affected by the impaired glutamine metabolism that we have demonstrated. Intriguingly, MYCN-regulated translation has recently been reported as a therapeutic vulnerability in medulloblastoma (69), and our studies support this possibility in high-risk neuroblastoma. Importantly, highly selective epitranscriptomic inhibitors are now emerging (70), and the interdependence of splicing and epitranscriptomic pathways suggests that drug combinations selectively targeting both pathways may be highly efficacious. Our work also highlights the interplay of arginine methyltransferases and RNA methylation, as previously suggested for PRMT1 and METTL14 (71).

PRMT5 inhibition using GSK3326593 led to a significant improvement in survival in the *Th-MYCN* neuroblastoma mouse model and excellent target engagement was observed with strong SDMA reduction. Pathways identified by our *in vitro* analysis as being downstream of the MYCN-PRMT5 axis, including splicing, epitranscriptomics, DNA damage and glutamine metabolism, were similarly altered. Cell death and tumour regression, however, were not apparent, but overall, our studies indicate that PRMT5 inhibition will be especially useful as part of combination therapies. Such studies are already yielding promising results in other cancers (72,73) and the detailed mechanistic analyses presented here will rationalize new actionable pathways for combination therapies of MYCN-driven cancers. Together with recent findings that MNA neuroblastoma is highly sensitive to indisulam (23,24), our study underlines the vulnerability of high-risk neuroblastoma to inhibitors of the spliceosome.

## MATERIALS AND METHODS

### Cell culture

Sources of neuroblastoma and immortalised disease-free cell lines used in this study are detailed in Supplementary File 2. LAN-1, LAN-5, Kelly, and GIMEN were cultured in RPMI 1640 (Gibco), and CHP-212, IMR32, NGP, SK-N-BE(2)C (BE2C), SHEP, SH-SY-5Y, SK-N-AS, SH-IN, NBL-S, LAN-6, RPE-1, NF-TERT, and NF-730 were cultured in Dulbecco’s modified eagle’s medium (DMEM):F12-HAM (Sigma). Both media were supplemented with 10% (v/v) FBS (Life technologies), 2mM L-glutamine, 100 U/mL penicillin, 0.1 mg/mL streptomycin, and 1% (v/v) non-essential amino acids. SH-EP-Tet21N (SHEP-21N) were cultured in RPMI 1640 (Gibco), supplemented with 10% (v/v) tetracycline-free FBS (Life technologies), 2 mM L-Glutamine, 100 U/mL penicillin, 0.1 mg/mL streptomycin, and 1 μg/mL tetracycline. Cell counts were measured using a Countess automated cell counter (Thermo Fisher Scientific). All cell lines were routinely tested for mycoplasma contamination and were confirmed to be mycoplasma negative. Main genotype details of cell lines are given in Supplementary File 2.

### Drug treatments, siRNA knockdowns, and cell proliferation

Prior to treatment with GSK3203591 (dissolved in DMSO), cells were seeded and allowed to adhere before being treated at the concentrations and durations indicated in figure legends in complete media. Control cells were treated with equivalent DMSO concentration. Transient siRNA knockdowns were performed by using short interfering RNA (Supplementary File 2). Reverse transfections were performed using 25 nM siRNA and Lipofectamine RNAiMAX (Invitrogen) complexed in OptiMEM media (Invitrogen) and added to cell suspensions prior to seeding and incubated for at least 48-hours before cell lysis. Cell proliferation was measured using the IncuCyte ZOOM live-cell analysis system with images taken every 4-hours.

### MTT Cell Viability and Colony Formation Assays

Cells were seeded in 96-well plates at a cell density ranging from 500-3000 cells/well and treated the next day in triplicate with a serial dilution of GSK3203591. After 120h, 10μL of MTT (5 mg/mL) (Sigma) was added/well, followed by 40μL of SDS lysis buffer (10% SDS (w/v), 1/2500 (v/v) 37% HCl) after 3 hours had elapsed. Following overnight incubation at 37°C, the plates were read at 570nm and 650nm, using SpectraMax 190 plate reader (Molecular Devices). For colony formation assays, cells were seeded into 6-well plates (1 × 10^3^ cells/well) and allowed to adhere for 24-hours before being treated with GSK3203591 or DMSO equivalent. Cells were re-dosed with GSK3203591 or DMSO every 96h. After 17-days cells were fixed in 4% paraformaldehyde and stained with methylene blue.

### Protein Extraction and Immunoblotting

Floating cells were collected and attached cells trypsinised and then combined and centrifuged at 4°C collected before being lysed in Radioimmunoprecipitation assay (RIPA) buffer with added protease and phosphatase inhibitors and sonicated on high for 30 seconds off 30 seconds on for 3 minutes (Diagenode Bioruptor). The protein concentration was determined using the Micro BCA TM protein assay kit (Thermo Fisher). Immunoblotting was performed as described previously (5) using antibodies in Supplementary File 2.

### RNA Extraction, Reverse Transcription, PCR assays

RNA was extracted from attached cells using QIAzol and the miRNeasy kit (QIAGEN) according to the manufacturer’s instructions, including DNAse treatment. cDNA synthesis was performed using Superscript IV cDNA synthesis kit (Invitrogen). Quantitative PCR (5 ng cDNA/well) was performed by using QuantiNova kit on Mx3500P PCR machine (Stratagene). Gene-specific primers were used for end-point PCR (HotStarTaq Plus DNA Polymerase; Qiagen), to detect inclusion or exclusion of alternative exons, after electrophoresis on agarose gels (2%). The oligonucleotide primers used to detect target gene expression and alternative splicing events are listed in Supplementary File 2.

### Cell Cycle Analysis

Cells were seeded and treated with GSK3203591 or DMSO equivalent for 24-96 hours. Cells were trypsinised, washed in PBS and dripped into ice cold 70% ethanol and kept at −20°C for at least 2-hours. Propidium-iodide labelling, and flow cytometry analysis was performed as previously described (5), and analysed using FlowJo software v10.

### RNA-seq and Bioinformatic Analysis

RNA was extracted as above and quantified using Nanodrop (ND1000) and quality confirmed using an Agilent ScreenTape RNA assay. Two biological replicates were used for RNA-seq of each condition with the paired-end option of 100bp reads on BGIseq-500 (BGI, Shenzhen, China). Differential gene expression bioinformatics and alignments were as previously described (74). Alternative splicing analysis was conducted with The Junction Usage Model (JUM) splicing analysis package (75). RNA-seq reads were aligned to the human genome (hg38) with the Spliced Transcripts Alignment to a Reference (STAR) protocol (76). We used a p=<0.05 and a ΔPSI >0.05 cut-off for assignment of differentially spliced genes (DSGs). Gene expression analyses of published neuroblastoma datasets and Kaplan-Meier analyses were performed by using the R2 Genomics Analysis and Visualization Platform (http://r2.amc.nl).

### Irradiation and Confocal Microscopy

Cells were seeded on rat tail collagen (10 ug/mL) coated coverslips 24-hours prior to treatment with GSK3203591/DMSO for 48 hours. Cells were then exposed to 2.5 Gy irradiation and incubated for 1-or 24-hours in media containing GSK3203591. Cells were fixed using 4% paraformaldehyde and washed 3x with PBS. Coverslips were permeabilised for 25 minutes in PBS containing 0.1% Triton-×100 (Merck, UK), blocked in PBS containing 10% (v/v) BSA + 0.1% (v/v) Triton-×100 (both Merck, UK) and primary antibodies (γH2AX, Cell Signalling Technologies, 1:200) diluted in antibody buffer (PBS containing 2% (v/v) BSA + 0.1% (v/v) Triton-X) were applied at 4°C overnight. Coverslips were washed 3x PBS and secondary antibodies (Alexa Fluor™ 555 anti-rabbit, A-21428, 1:1000) applied in antibody buffer for 1-hour in the dark at room temperature, washed with PBS and incubated 5 minutes with PBS containing 2 µg/mL Hoechst (33342, Fisher Scientific, UK) and washed in PBS and briefly in ddH2O then mounted on glass slides using ProLong GoldTM anti-fade glass mounting medium (P36980, Fisher Scientific, UK). Z-stack images were acquired using a Leica SP8 confocal microscope at magnifications indicated in legends using 2048×2048 pixel resolution. Foci per nucleus were quantified using CellProfiler 4.2.6 using maximal orthogonal projection of Z-stacks.

### m6A-Methylated RNA immunoprecipitation (MeRIP)-qPCR

Cells were treated with GSK3203591/DMSO equivalent for 96 hours prior to being trypsinised and collected. Total RNA was prepared using RNeasy Midi kit (Cat# 75144, QIAgen, UK) as per manufacturer’s instructions and DNase digest was performed on column using RNase free DNase set (Cat# 79254, QIAgen, UK). RNA was quantified using NanoDrop1000 and 75 ug total RNA was used to purify mRNA using Dynabeads® mRNA Purification Kit (Cat# 61006, Fisher, UK) and RNA clean up performed using RNeasy MinElute Cleanup Kit (Cat# 74204) both as per manufacturer’s instructions. ND-Methyladenosine RNA immunoprecipitation (meRIP) was performed using Magna MeRIP™ m6A Kit (Cat# 17-10499, Sigma-Aldrich, UK) as per manufacturer’s instructions. Briefly, protein A/G beads were washed in 1x IP buffer three times and coupled to 5 µg m6A antibody (MABE1006) or IgG (CS100621) at room temperature for 30-minutes. Beads were then washed three times and resuspended in meRIP reaction mixtures containing 0.5 µg unfragmented in-tact mRNA and 1% (v/v) RNase inhibitor in 1x IP-buffer and incubated with rotation for 2 hours at 4°C. Beads were washed three times in IP buffer and transferred to a clean tube. Immunoprecipitated transcripts were eluted by resuspending beads in elution buffer containing 6.67 mM ND-Methyladenosine, 5’-monophosphate sodium salt (CS220007) and RNase inhibitor in 1x IP buffer and incubated for 1 hour with shaking at 4°C. Eluates were transferred to clean tubes and RNA clean up performed using RNeasy MinElute Cleanup Kit (Cat# 74204) as per manufacturer’s instructions. cDNA was synthesised as above on 10% (50 ng) mRNA input, m6A-IP and IgG-IP samples and then analysed using RT-qPCR, as above.

### Stable isotope labelling

Cells were seeded 24-hours prior to treatment with 2.5 µM GSK3203591 for 72-hours before being washed with PBS and incubated for 8-hours with DMEM media supplemented with 10% dialysed FBS, 1% pen/strep, 10 mM glucose (unlabelled or ^13^C_6_-glucose labelled), 2 mM glutamine (unlabelled or ^13^C_5_-glutamine labelled). Media was removed and cells washed twice in ice-cold saline. Plates were then placed on dry ice and scraped into 800 µL ice-cold 80% LC-MS grade methanol, centrifuged at 15,000 x g for 10 minutes 4°C and stored at −80°C.

Cellular metabolites were extracted and analysed by gas chromatography-mass spectrometry (GC-MS) using protocols described previously (77). Metabolite extracts were derived using N-(tert-butyldimethylsilyl)-N-methyltrifluoroacetamide (MTBSTFA) as described previously (78). An internal standard, D-myristic acid (750ng/sample), was added to metabolite extracts. Mass isotopomer distribution was determined using a custom algorithm developed at McGill University (77).

### SunSet Assay

Translation of nascent protein was measured using the previously described SUrface SEnsing of Translation (SUnSET) assay (79). Cells were seeded 24-hours prior to treatment with GSK3203591/DMSO for 72h then pulsed with 1.25 µM puromycin for 1-hour prior to cell lysis and protein extraction (as above). Protein extracts of puromycin pulsed and non-puromycin pulsed control cells were subject to immunoblot (as above) using anti-puromycin antibody (clone 12D10, 1:1000, Sigma).

### In vivo evaluation of GSK3326593 in Th-MYCN GEMM mice

All experiments were approved by The Institute of Cancer Research Animal Welfare and Ethical Review Body and performed in accordance with the UK Home Office Animals (Scientific Procedures) Act 1986, the United Kingdom National Cancer Research Institute guidelines for the welfare of animals in cancer research and the ARRIVE (animal research: reporting *in vivo* experiments) guidelines. The study was performed using both male and female hemizygous mice, which developed palpable tumours at 50 to 130 days with a 25% penetrance. Transgenic Th-*MYCN* mice were genotyped to detect the presence of human *MYCN* transgene. Tumour development was monitored weekly by palpation by an experienced animal technician. Mice with palpable tumours at >3 mm were treated with either GSK3326593 at 100mg/kg, twice per day by oral gavage or vehicle (0.5% methylcellulose) twice per day by oral gavage. Mice were housed in specific pathogen-free rooms in autoclaved, aseptic microisolator cages (maximum of four mice per cage). Mice were allowed access to sterile food and water ad libitum.

### Statistical Tests

The normality of datasets ≥3 datapoints was determined using a Shapiro-Wilks test. Normally distributed datasets were analysed using a t-test with a significant p-value of 0.05 and data were plotted using GraphPad Prism 9.4.0. Statistical tests, sample size and the number of biological replicates performed for specific experiments are detailed in figure legends.

## DATA AVAILABILITY

RNA sequencing data is available from the European Nucleotide Archive (ENA) accession PRJEB72851 / ERP157650.

## ACKNOWLEDGEMENTS

We are very grateful to Professor Vande Velde (Universite de Montreal, Canada) for the kind gift of anti-hnRNPA1B antibodies. We would like to thank D. Avizonis and L. Choinière from the McGill University Metabolomics Core Facility for their support. We would like to acknowledge Yann Jamin and Barbara Martins da Costa for the help with animal work. The authors gratefully acknowledge all members of the Wolfson Bioimaging Facility, the Flow Cytometry Facility and the Bristol Genomics Facility at The University of Bristol for their support and assistance in this work. We would also like to thank Dr Andy Fedoriw and Dr Jan Kosters for feedback and helpful comments. Ipsen reviewed this manuscript for scientific accuracy, but had no input into the content. We thank Dr Heather Etchevers for human neural crest cell line RNA.

## AUTHOR CONTRIBUTIONS

KM and LC conceived the study and co-ordinated the project. JoB, MK, MS, KM designed the experiments and with JB, AM, SM, AG, JH-P conducted the experiments with technical help from KG. JoB, MS, AP and KM performed bioinformatics. MK, MS, DL, NJ and EV conducted metabolomics experiments. JoB, MK, MS, JB, DL, EV, EP, LC and KM collected and analysed data. EP carried out mouse experiments and analysed with LC. LC was responsible for and supervised the mouse studies. MK and KM drafted the manuscript. MK, MS, EP, AM, EV, AG and NC helped to further the draft manuscript. JoB and KM wrote the final manuscript. All co-authors read and gave input to help improve the manuscript. Jo.B, M.K, M.S are considered equal first authors and listed alphabetically and agreed that each may list themselves first in their individual documentation.

## SUPPLEMENTARY FIGURE LEGENDS

**Supplementary Figure 1:**
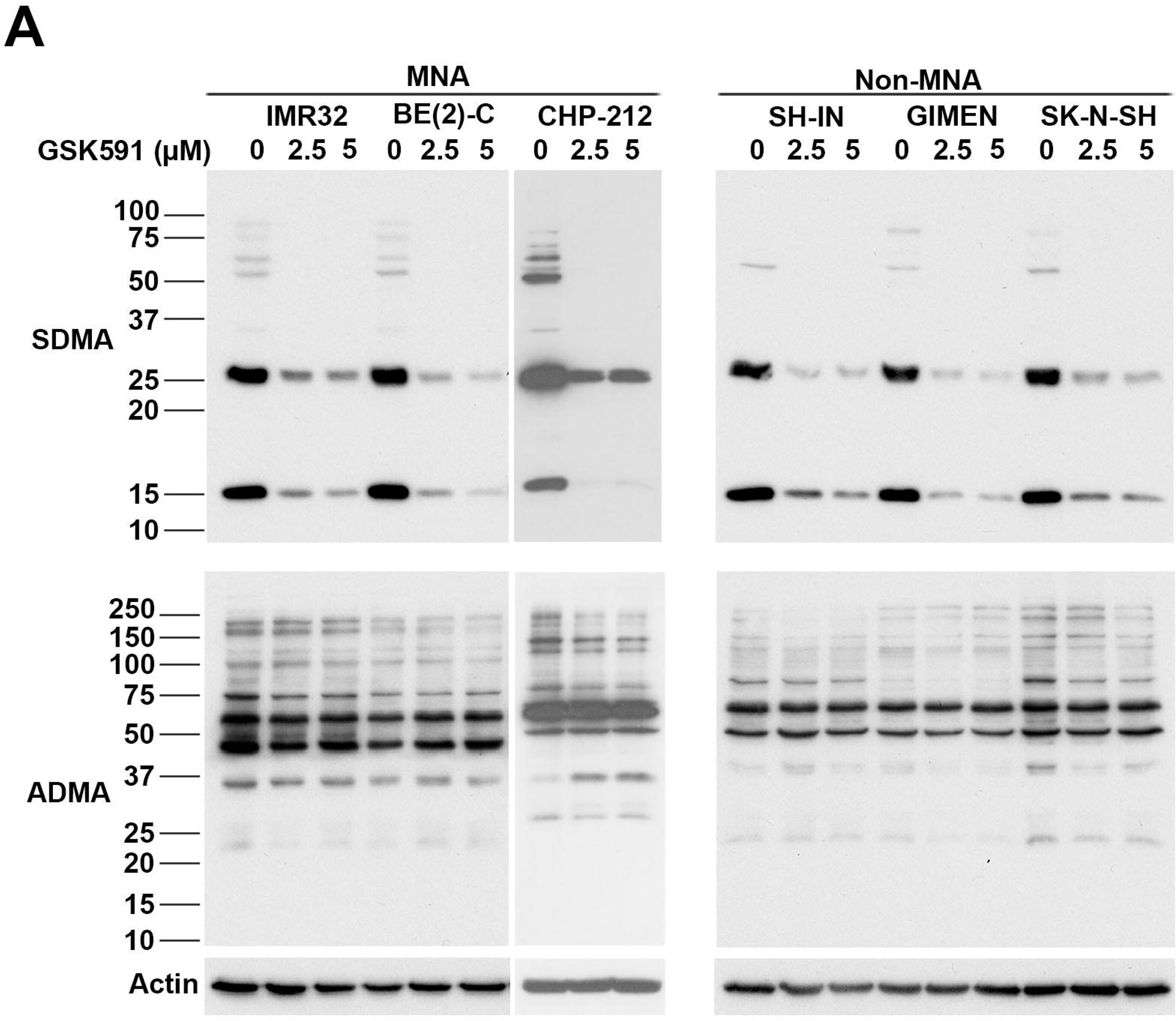
**[A]** Immunoblot of MNA and non-MNA cell extracts probed with anti-SDMA or anti-ADMA antibodies demonstrating on-target SDMA reduction following 0 (DMSO), 2.5, 5 μM GSK3203591 treatment. One representative experiment of 2 shown.

**Supplementary Figure 2:**
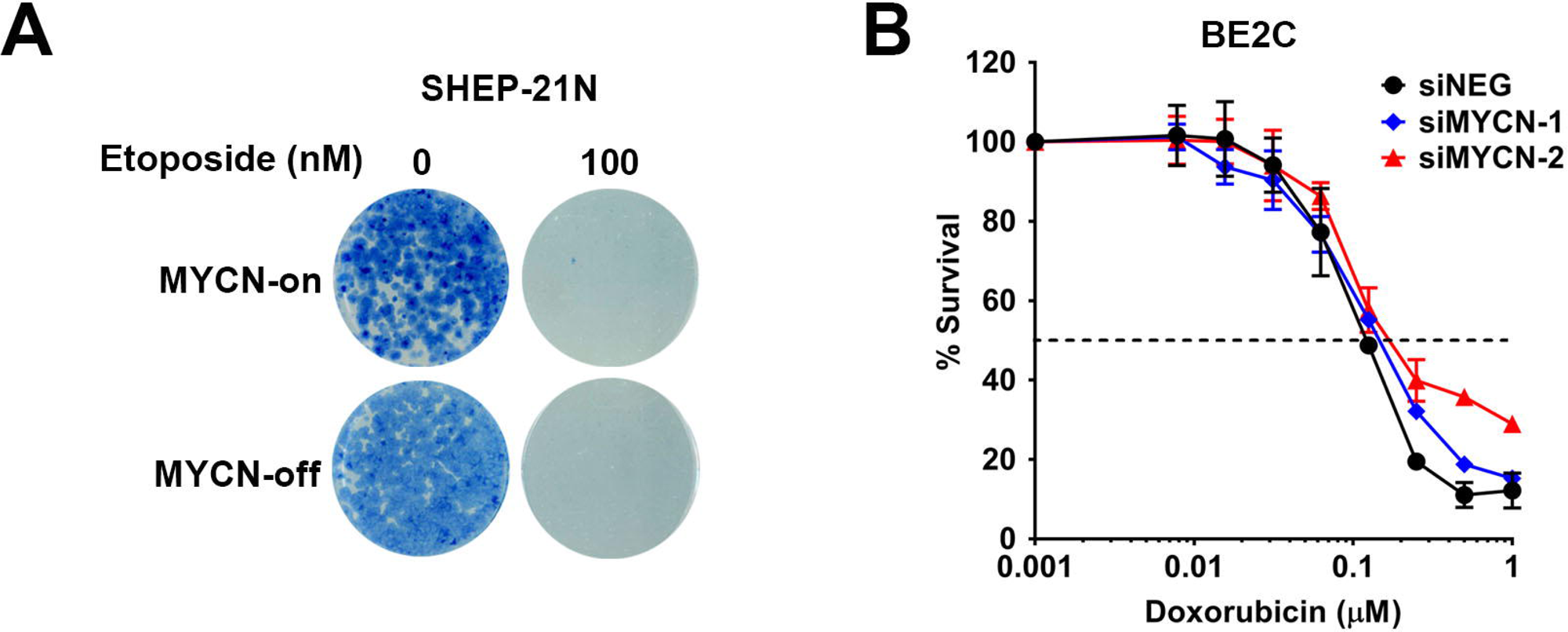
**[A]** Clonogenic assay of SHEP21N cells incubated with 0 or 100 nM etoposide with (MYCN off) or without (MYCN on) tetracycline showing MYCN-independent activity of Etoposide. One representative experiment of 2 shown. **[B]** Cell survival assay (MTT) of BE2C cells transfected with two different siRNA’s targeting MYCN or siNEG and incubated with increasing concentrations of Doxorubicin for 72hrs (n=3, mean ± SD).

**Supplementary Figure 3:**
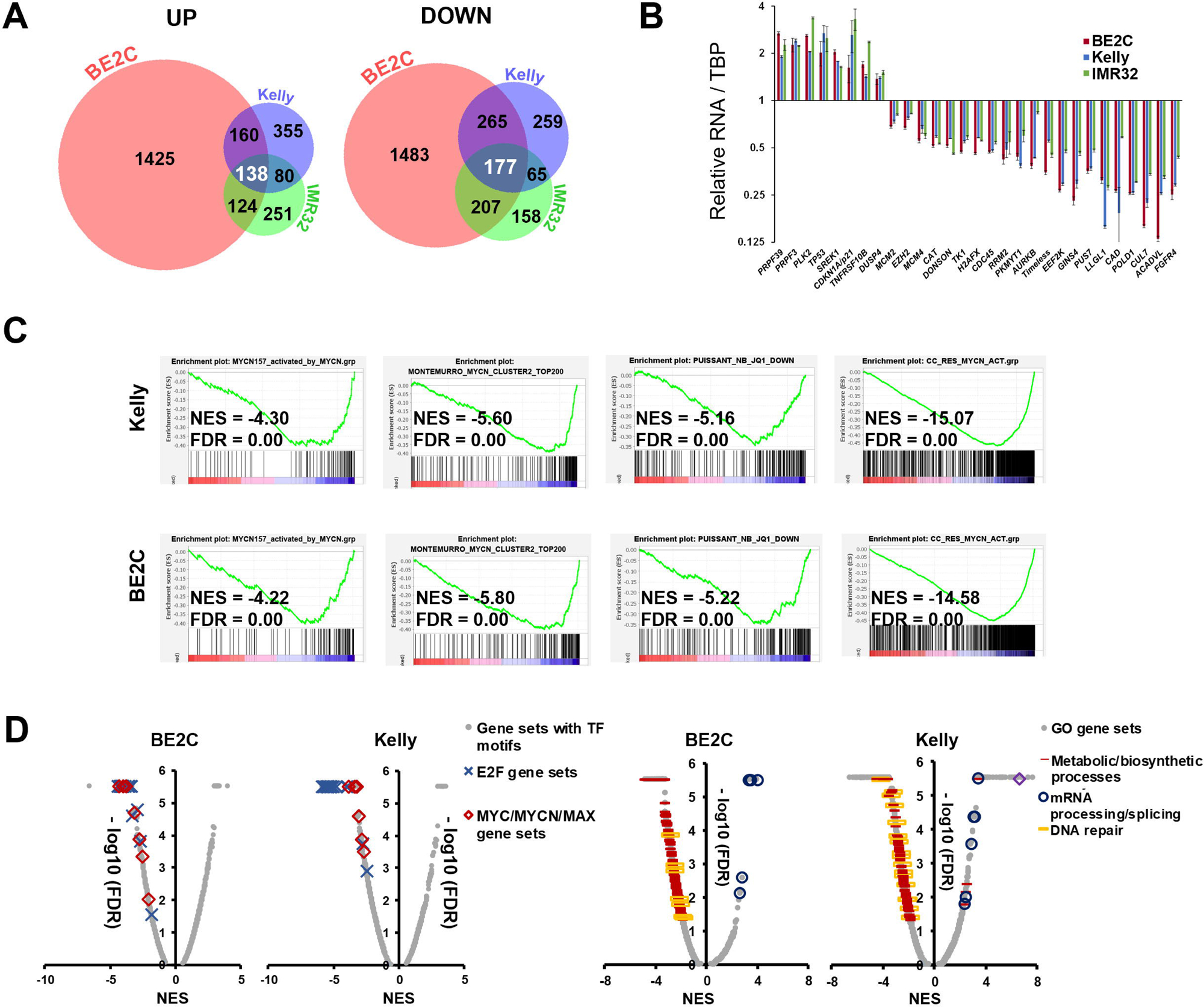
**[A]** Venn diagram of differentially expressed genes upregulated (left) or downregulated (right) following GSK3203591 treatment in 3 MNA cell lines. **[B]** RT-qPCR validation of 29 differentially expressed genes identified by RNAseq in MNA cell lines (n=2, mean ± SEM). **[C]** GSEA analysis of GSK3203591-treated BE2C (left) and Kelly (right) cells demonstrating strong inhibition of MYCN-dependent gene sets. **[D]** Global summary of GSEA plots showing repression of MYC/MYCN and E2F gene sets (left) and other frequently altered gene ontology GSEAs affected by PRMT5 inhibition (right) in BE2C and Kelly cells treated with GSK3203591. NES, normalised enrichment score; FDR, false discovery rate.

**Supplementary Figure 4:**
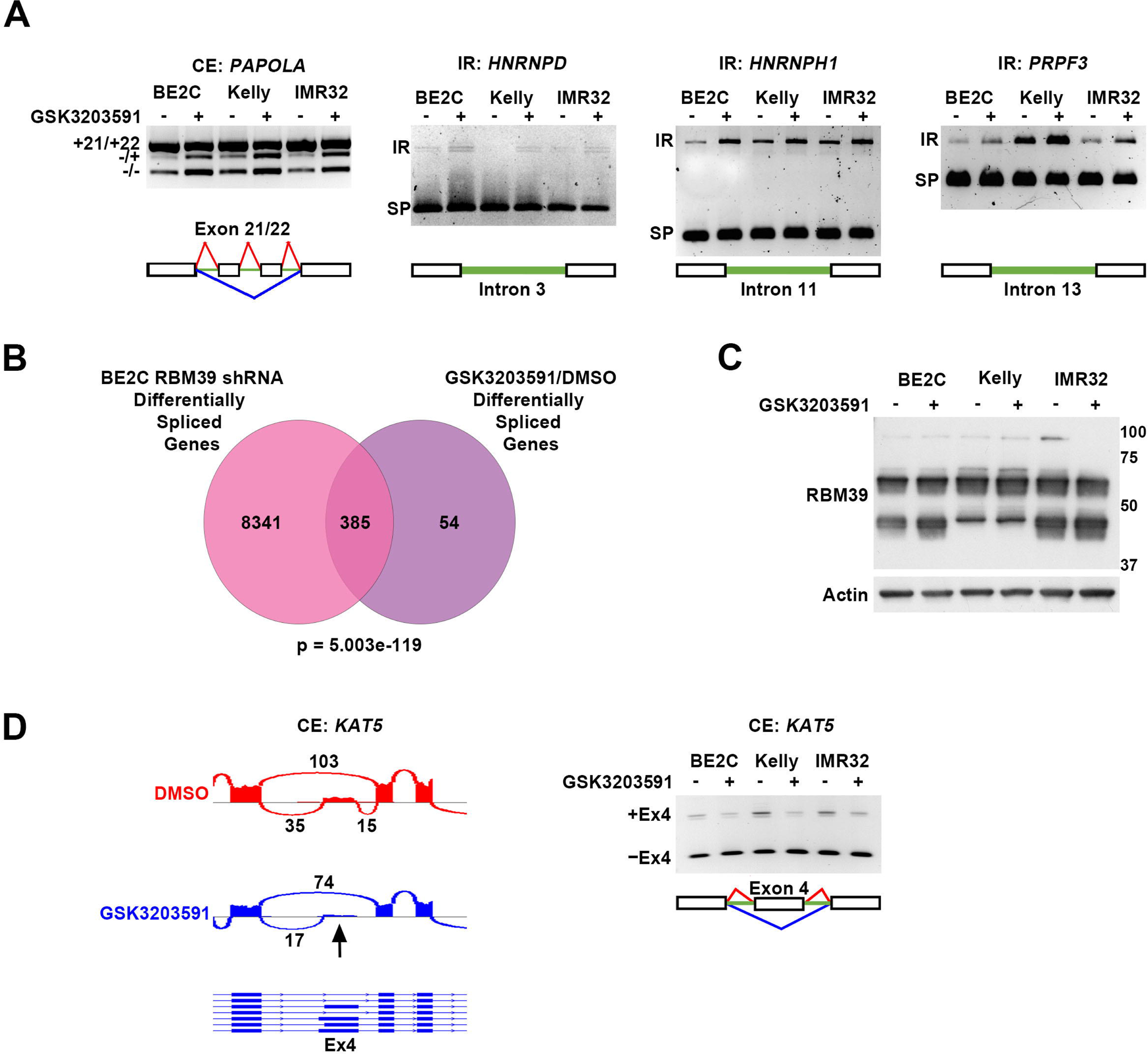
**[A]** End-point PCR validation of alternative splicing events in MNA cell lines treated with GSK3203591. One representative experiment of 3 shown. **[B]** Venn diagram of alternatively spliced genes in BE2C cells transfected with two separate shRNA’s targeting RBM39 and alternatively spliced genes shared by the 3 neuroblastoma cell lines following GSK3203591 treatment in this study. **[C]** Immunoblot (IB) of MNA cell lines treated with GSK3203591 showing no change to RBM39 expression. One representative experiment of 2 shown. **[D]** Representative sashimi plot (left) and end-point PCR validation (right) of *KAT5* cassette exon alternative splicing event after GSK3203591 showing decreased inclusion of exon 4. One representative end-point PCR experiment of 2 shown.

**Supplementary Figure 5:**
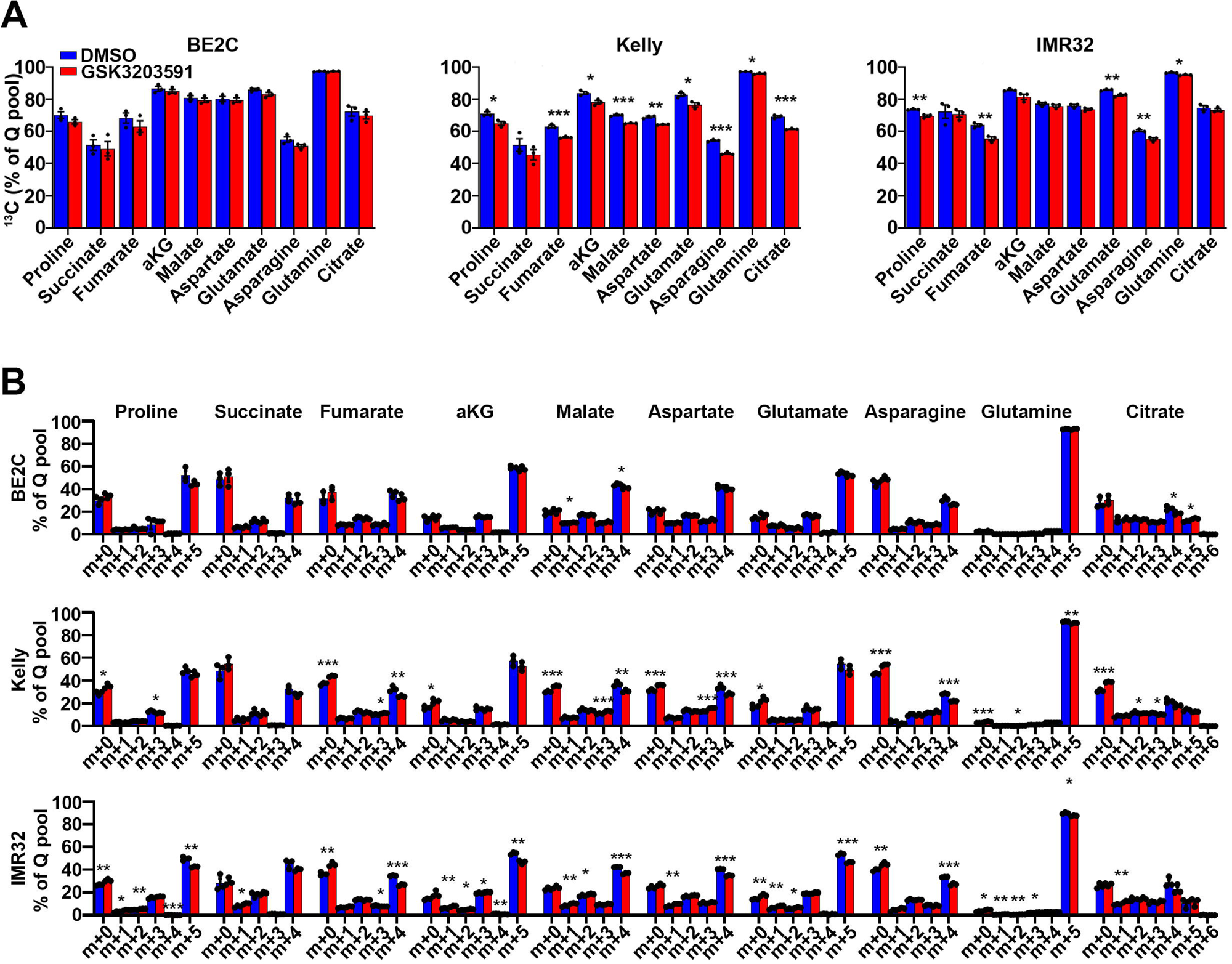
**[A]** Percentage of ^13^C labelling in metabolite pools from U-[13C]-Q. Asterisks denote the p-value obtained using t-tests (* p<0.05, ** p<0.01, *** p<0.001). **[B]** Mass isotopomer distribution (MID) analysis for U-[13C]-Q labelling in neuroblastoma cell lines for Proline, Succinate, Fumarate, αKG, Malate, Aspartate, Glutamate, Asparagine, Glutamine, and Citrate.

**Supplementary Figure 6:**
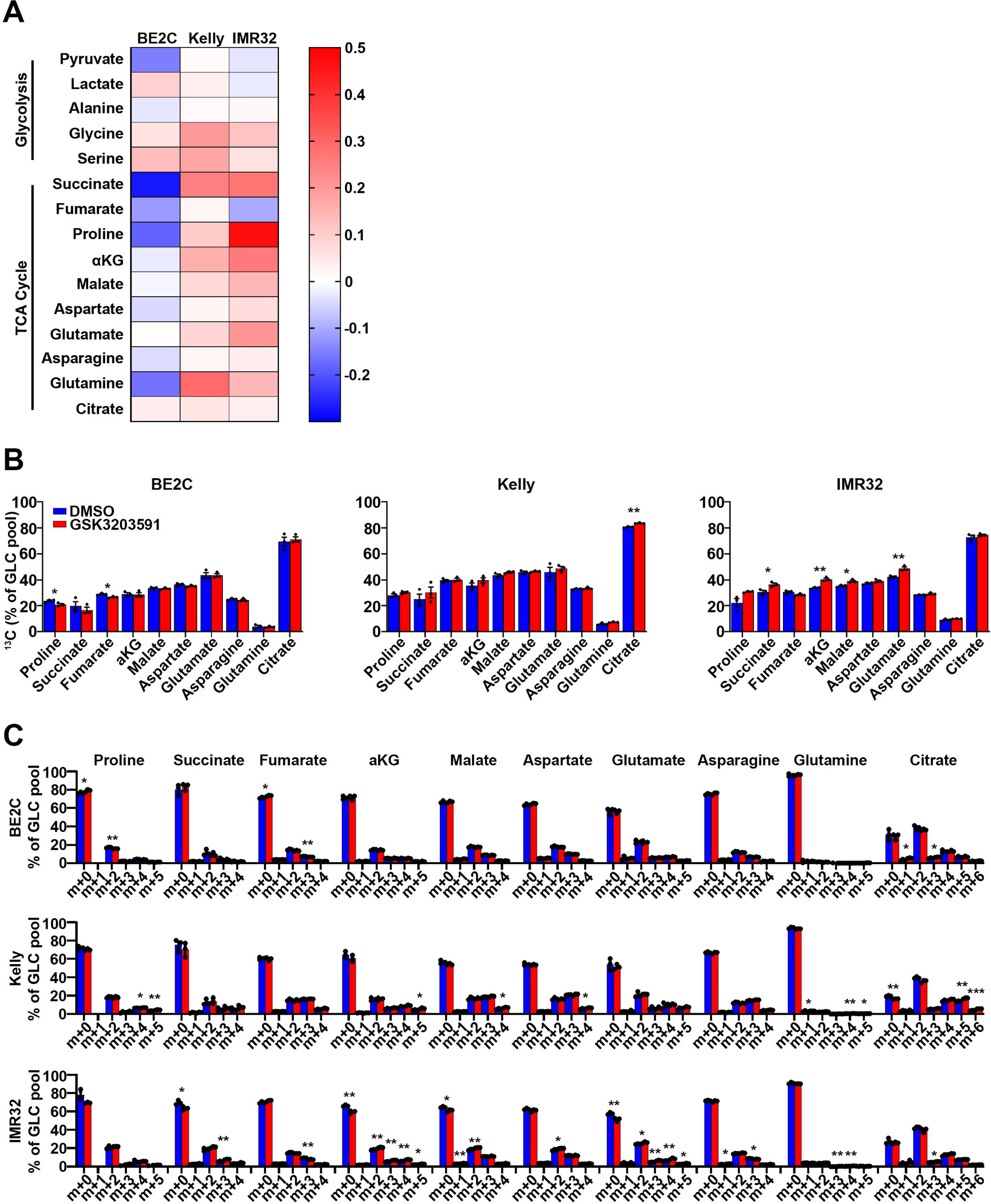
**[A]** Heatmap summarizing glucose incorporation into TCA cycle intermediates after PRMT5 inhibition in three cell lines (BE2C, IMR32 and Kelly), data derived from stable isotope labelling experiments (data represented as the average of 3 independent experiments). **[B]** Percentage of ^13^C labelling in metabolite pools from U-[13C]-GLC. Asterisks denote the p-value obtained using t-tests (* p<0.05, ** p<0.01, *** p<0.001). **[C]** Mass isotopomer distribution (MID) analysis for U-[13C]-GLC labelling in neuroblastoma cell lines for Proline, Succinate, Fumarate, αKG, Malate, Aspartate, Glutamate, Asparagine, Glutamine, and Citrate.

**Supplementary Figure 7:**
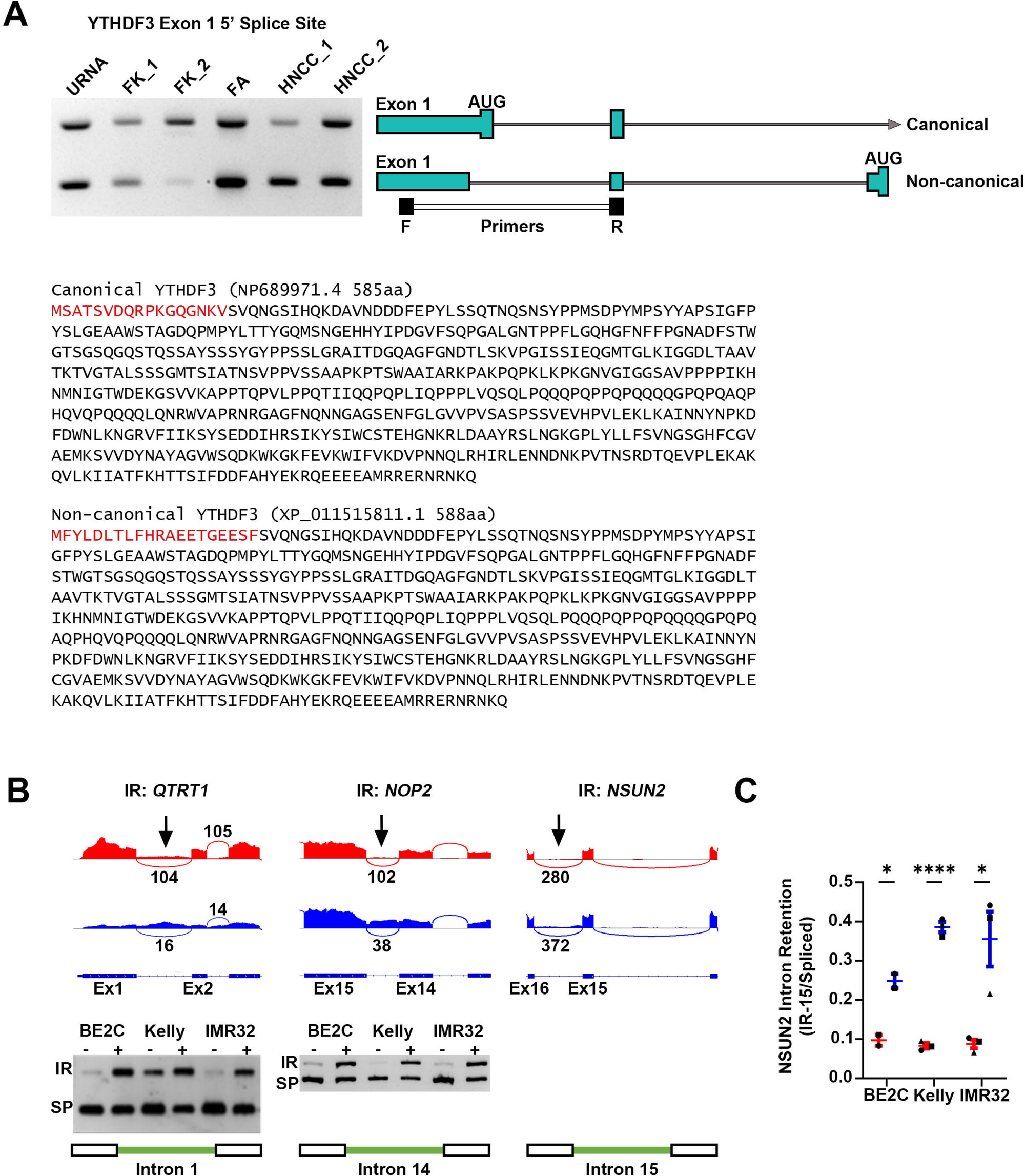
**[A]** YTHDF3 5’-alternate splicing in *YTHDF3* exon 1. End point PCR shows usage of both isoforms in fetal kidney (FK), universal RNA (URNA), fetal adrenal (FA) and human neural crest cells (HNCC) (left). Schematic demonstrate alternative 5’ splice site in exon 1 (right). Annealing sites of primers in exon 1 (forward, F) and exon 2 (reverse, R) produce a larger PCR product when the canonical isoform of YTHDF3 is expressed. Protein sequences (below) demonstrates YTHDF3 proteins based on exon 1 (canonical) and exon 3 (non-canonical) AUG usage. **[B]** Representative sashimi plot (top) and end-point PCR validation (bottom) of alternative splicing events in RNA modifier proteins QTRT1, NOP2 and NSUN2 following GSK3203591. IR, intron retention; SP, spliced product). **[C]** RT-qPCR validation of intron retention of intron 15 in *NSUN2* following GSK3203591 (data relative to TBP, n=
≥2, mean ± SEM, unpaired t-test, *=p<0.05, ****=p<0.0001).

